# Mechanistic Studies of the Stabilization of Insulin Helical Structure by Coomassie Brilliant Blue

**DOI:** 10.1101/2020.08.26.267799

**Authors:** Sandip Dolui, Ranit Pariary, Achintya Saha, Bhisma N Ratha, Amaravadhi Harikishore, Susmita Saha, Snehasikta Swarnakar, Anirban Bhunia, Nakul C Maiti

**Author notes:** both contributed equally (Sandip Dolui, Ranit Pariary). Department of Phytopharmaceuticals, Centurion University of Technology and Management, Alluri Nagar, Parlakhemundi 761211, India (Bhisma N Ratha). Address correspondence to Nakul C. Maiti, Division of Structural Biology and Bioinformatics, CSIR-Indian Institute of Chemical Biology, 4, Raja S.C. Mullick Road, Kolkata 700032, India. Anirban Bhunia, Department of Biophysics, Bose Institute, P-1/12 CIT Scheme VII (M), Kolkata 700054, India.

## Abstract

Human insulin (HI) is an essential protein hormone and its biological activity mostly depends on folded and active conformation in the monomeric state. The present investigation established that Coomassie Brilliant Blue G-250 (CBBG), a small multicyclic hydroxyl compound can reversibly bind to the hormonal protein dimer and maintained most of α-helical folds crucial for biological function of the enzyme. The solution-state 1D NMR and isothermal calorimetric analysis showed a sub-micromolar binding affinity of the molecule to HI. 2D NOESY NMR established that the HI dimer undergoes residue level local conformational change upon binding to CBBG. The chemical shift perturbation and the NOE parameters of active protons of amino acid residues throughout the polypeptides further suggested that CBBG upon binding the protein stabilize α-helixes of both the A and B subunits of the hormonal protein. The changes in Gibb’s free energy (∆*G*) of the binding was of ~−11.1 kcal/mol and suggested a thermodynamically favourable process. The changes in enthalpy (∆*H*) and entropy term (*T*∆*S*) were −57.2 kcal/mol and −46.1 kcal/mol, respectively. The negative changes in entropy and the NOE transfer effectiveness of several residues in the presence of CBBG molecules indicated that the binding was an enthalpy driven favourable equilibrium process. The NMR-based atomic resolution data and molecular docking studies confirmed that the CBBG binds to HI at the dimeric stage and prevents the availability of the crucial residue segments that partake directly in further oligomerization and subsequent fibrillation. Extended computational analysis based on chemical shift perturbation of protons of active residues further established receptor-ligand based pharmacophore model comprised of 5 hydrophobic and a hydrogen bond acceptor features that can anchor the residues at the A and B chains of HI and inhibit the partial unfolding and hydrophobic collapse to nucleate the fibrillation. Taken together, the results demonstrated that CBBG and their close analogues might be useful to develop a formulation that will maintain the active and functional form of the hormonal protein for a significantly longer time.

**TOC:** 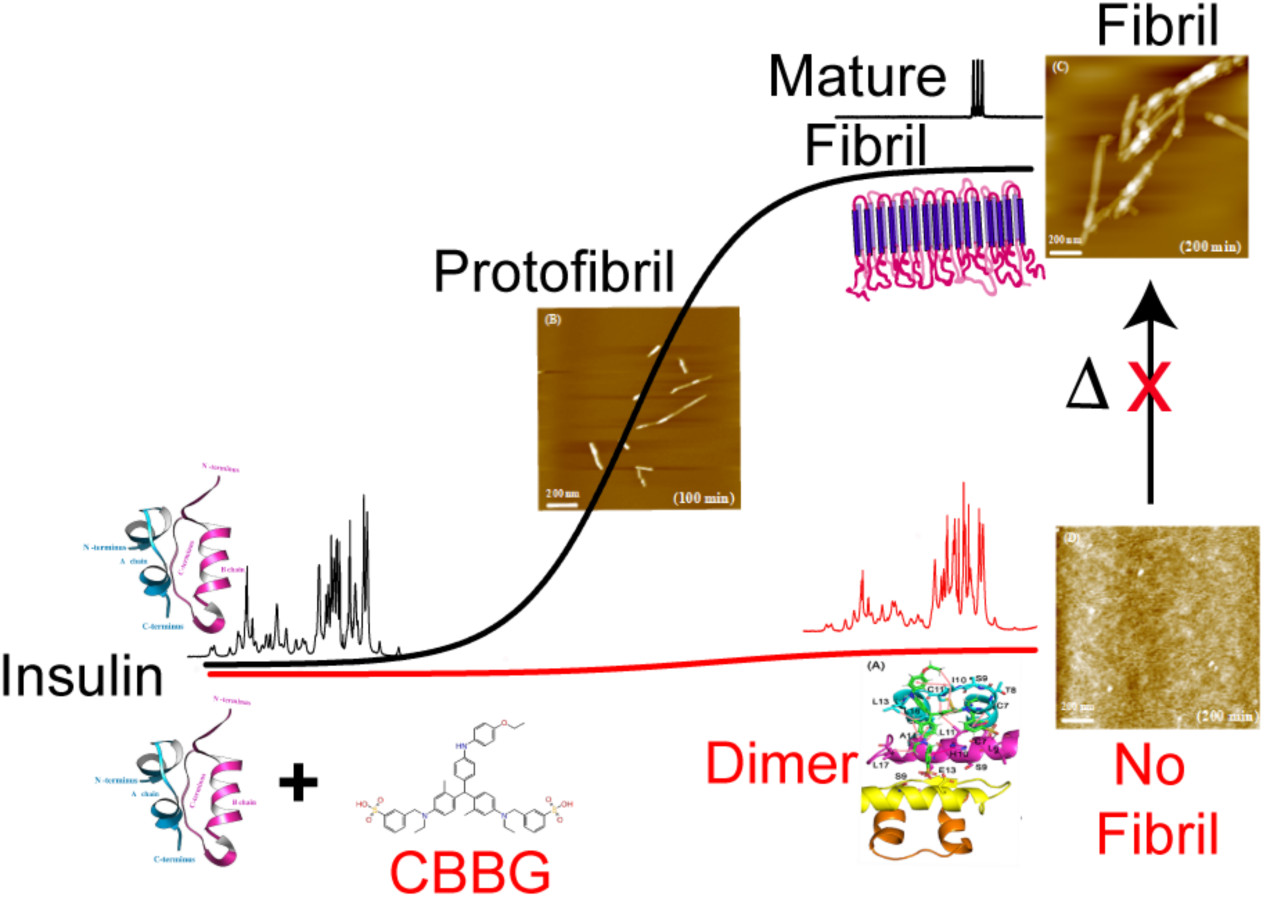

## Introduction

Human insulin (HI) is a 51 residues long protein hormone produced in the pancreas and it plays a significant role in glucose metabolism and helped to maintain the normal glucose level in blood.^1–5^ After its synthesis, HI is stored in the pancreatic β-cells in a hexameric assembly state.^6–8^ HI, in its monomeric form, was comprised of an A chain (21 amino acids) and a long B chain (30 amino acids) linked together by a pair of inter-chain disulfide bonds (Figure 1A).^9–11^ Structurally, A-chain was composed of two α-helices at its N-terminus (A_2–8_) / C-terminus (A_13–19_) which are interconnected by an intra chain disulfide linkage forming a short loop at its N-terminus (Figure 1B). In contrast, the B-chain (in its monomer form) was mainly with α-helical central region (B_8–19_) flanked by extended N/C terminal segments (Figure 1B). In contrast to A chain, the terminal sequences of HI B chain are shown to adopt extended helical conformation in the presence of allosteric ligands.^12,13^ Strikingly, presence of cofactors (Zn^++^) and their coordination with histidine (His^B10^) residues was sufficient to induce structural transition from monomer into hexamer state.^14,15^ As a consequence, the N-terminus (B_1-8_) of B chain was shown to adopt two possible states in response to their allosteric ligand concentrations. Therefore, the N-terminus residues of the B-chain adopted an extended conformation as a tense state (T)^12^ in absence of ligands or low Cl^−^ ions or, a helical relaxed state (R) conformation^16^ in presence of allosteric ligands or high Cl^−^ ions, respectively. This kind of conformational transitions and partial unfolding of the protein was also evidenced in the process of its receptor binding.^12,17^

**Figure 1.**
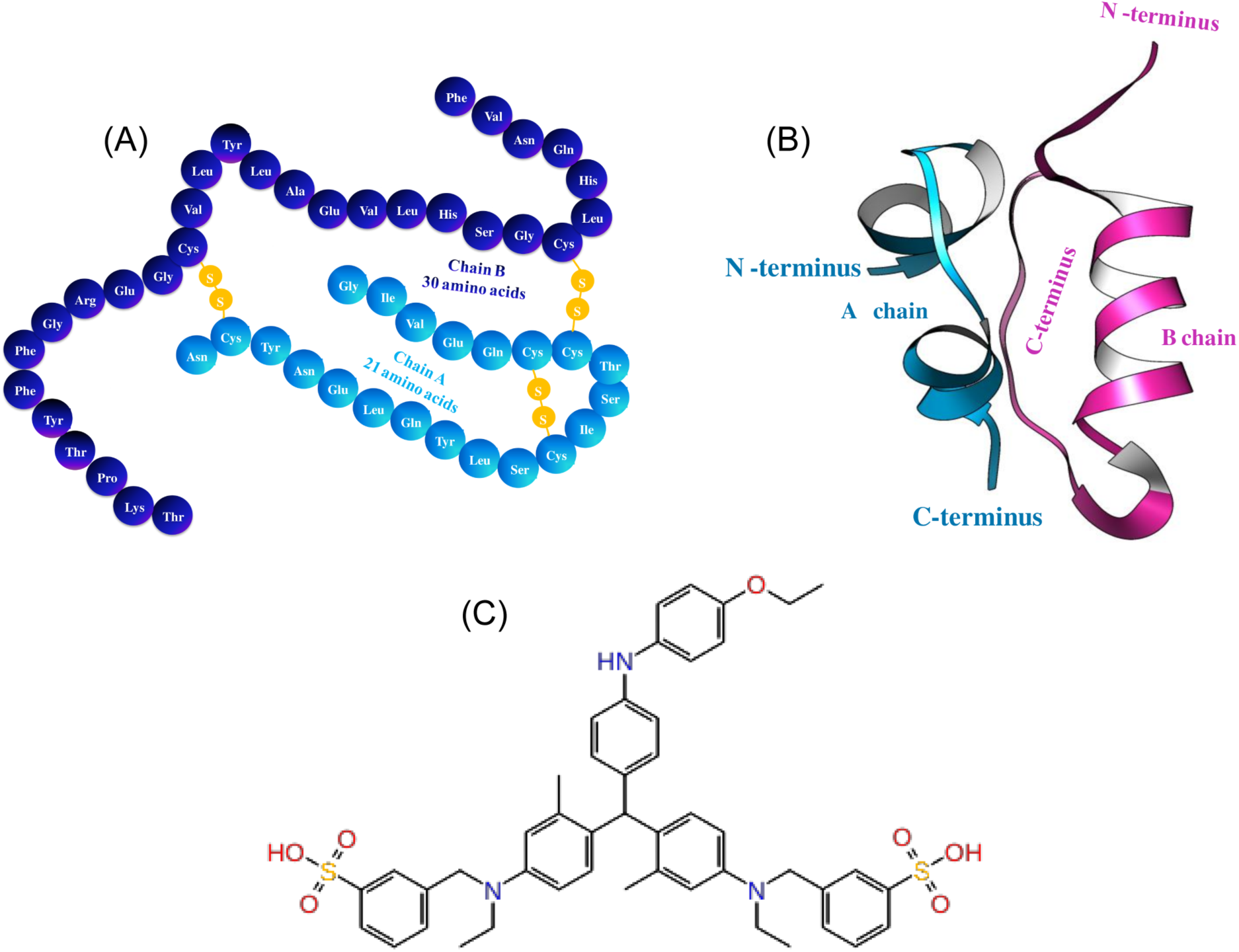
Structure of human insulin (HI) and Coomassie Brilliant Blue G-250 (CBBG). (A) the primary structure of human insulin. The A-chain (skyblue) consists of 21 amino acids and B-chain (deep blue) consist of 30 amino acids they linked by two interchain and one intrachain-disulphidebridge that are highlighted in yellow. (B) The left panel shows a secondary structure of insulin (RCSB Protein Data Bank (PDB) entry 1GUJ; illustrated in Pymol). The A-chain (skyblue) consists of two α-helices. The B-chain (pink) features an α-helix and a random coil. (C) The lower panel shows the stick model of Coomassie Brilliant Blue G-250 (CBBG).

Once released into blood circulation, HI molecules are exposed to undergo dynamic structural transitions from a hexameric form to biologically active monomer^18^ in response to changes in its concentration, physiological pH as well as allosteric ligands. One of the key determinants to maintain structural stability and function of the HI molecule is to maintain the integrity of helical folds during structural transitions and unfolding steps of HI B chain. This innate loss of structural stability and compactness of the HI structure was increasingly recognised to be associated with a pathological state referred to as amyloidoma in diabetes patients.^15,19^ In this pathological conditions, HI molecule acquires the propensity to form amyloid like fibril aggregates at the site of HI injection. Furthermore, additional factors such as low pH/high temperature were also suggested to promote the aggregation and fibril formation^9,14,15,20–23^ and affect the stability of HI molecule and thereby reducing its shelf life.^21,24^

In the event of fibril formation, it is reported that the α-helical domains of the protein open up its hydrophobic regions and in the subsequent event of their collapse triggers amyloidogenesis and formation of unwanted fibrillar aggregates.^25–27^ This physicochemical process often proceeds with an elongated lag-phase, and the subsequent formation of small oligomeric intermediates followed by a sudden growth of formation of β-sheet rich fibrillar aggregates.^14,15,20–23,28–33^ Using different human HI mutant forms, Nielsen et al proposed a model for HI fibril formation and suggested that a partially folded intermediate triggers the hydrophobic collapse of two monomeric units. This collapse serves as an important starting point for the fibrillation of the protein both in the acidic and physiological pH condition.^34^ Using solution-state NMR spectroscopic analysis, Hua et al reported that in the early event of fibrillation, partial unfolding of N-terminal helixes of both the A and B chain occurs and the chains detach at least partially from the core of the protein.^27^ They further suggested that the novel detachment of N-terminal helical segments makes possible aberrant hydrophobic (protein/protein) interactions that create the amyloidogenic nucleus and eventually transformed into β-sheet rich fibrillar aggregates.

To avoid unwanted loss of the protein due to fibrillation, the pharmaceutical formulation of the protein hormone is often made under hexamer stabilizing conditions at near-neutral pH adding zinc and phenolic excipients.^35–37^Analogues of the protein are also made to overcome this effect, however, some of the analogues also showed an increased level fibril formation propensity.^21,24,35,38,39^ In the current investigation, we examined the binding interaction of Coomassie Brilliant Blue G-250 (CBBG) with HI and, its stabilizing effect on the helical folds present in the two peptide chains (A and B) of the protein. Using low- and high-resolution spectroscopic techniques such as circular dichroism (CD) and NMR spectroscopies, respectively, our investigation showed residue specific interaction of the protein and the molecule; it confirmed that the CBBG binding can arrest the α-helical folds of the protein even at an amyloidogenic condition (T = 60 °C, 25 mM HCl, 100 mM NaCl, pH ~1.6) and strongly inhibit the protein fibrillation. The CD spectral analysis demonstrated retention of the native α-helix for a longer period of time in the presence of CBBG. The isothermal calorimetric analysis provided several binding parameters and suggested that the stabilization processes was enthalpy driven. Together, our results highlighted that the molecule can provide strong thermal stability to HI and could restrict its partial unfolding event, even at an elevated temperature. The most intriguing and important observation, however, was that the residues which produced the partial fold to nucleate the fibrillation are indeed trapped by CBBG and provided significant stability to the helical domains of the hormonal protein. Further, a moderate binding affinity and high thermal (enthalpy) stability of CBBG/HI interaction strongly suggest that the molecule may be potentially useful in the formulation of HI and used as drug-adjuvant/excipient. In addition, the synergic investigation by NMR and computational analysis established a pharmacophore-model that may aid in designing other small molecules as the stabilizing agent for this wonderful hormonal protein, insulin.

## Results

### Non-toxic CBBG stabilizes the α-helical conformation of HI

Human insulin (HI) is a globular protein hormone and it contains three helical folds distributed in the A and B polypeptide chains of the protein (Figure 1B). In a harsh condition (T = 60 °C, 25 mM HCl, 100 mM NaCl, pH ~1.6) it produces a ‘novel partial fold’ in which the N-terminal segments of both the A- and B-chains partially detach from the core and one of the α-helices of the A-chain nucleate the fibrillation process. The present investigation revealed how a non-toxic small molecule, CBBG (Figure 1 and Supporting Figure S1) can retard this unfolding and protect the helical folds even in this severe solution condition of high temperature and low pH. To realize the stabilization effect of the molecule on the protein’s secondary structure, far UV circular dichroism (CD) spectroscopic measurements were carried out in the presence and absence of CBBG in acidic condition. Several panels in Figure 2 display the far UV CD spectra of HI in different solution conditions. The circular dichroic (CD) spectrum of HI in its initial state (before heating) showed two minima at ~ 208 and ~ 222 nm (black curves in Figure 2A and 2B, upper panels), suggestive of predominant presence of α-helical structure and in consonance with observations made by previous investigators.^10,43^ As the incubation continued at 60 °C, pH ~1.6, the ellipticity at the two minima was attenuated and a single minimum appeared at around 218 nm (Figures 2A and 2B, upper panels). This suggested that HI converted from mostly α-helix to a β-sheet structure (Figures 2A and 2B, lower panels) and, similar events were reported earlier.^10,43^ This transformation of the protein’s secondary structure to a cross β-sheet conformation is the good signature of fibrillation of HI. When CBBG was added to HI solution in the molar ratio of 1:1, it produced a CD spectrum similar to free HI at 0 h of incubation (red curves in the upper panel of Figure 2A). and, thus the major structural component was α-helical (lower panel of Figure 2A). Interestingly, α-helical CD signature of HI was largely preserved even after heating at 60 °C for 2 days (Figure 2A, shows the 36 h spectrum). The ratio of the ellipticities at 222 nm and 208 nm for the incubated protein in the presence of CBBG was similar to HI solution at 0 h of incubation and, it remained almost unchanged in the long incubation period. The bottom panel in figure 2A shows minimal conformational changes occurred in the presence of CBBG. These results suggested that CBBG can stabilize the native-like solution structure of HI and retarded the α-helix to β-sheet conformation transition. Upper panel in Figure 2B shows the CD spectrum obtained at different time point of incubation of the protein at the same conditions except the protein concentration was kept quite high (320 μM). The CD signatures (Figure 2B) of this sample also indicated the formation of β-sheet structure and, due to large amount of fibril formation, background scattering contribution made the baseline erroneous. However, the scattering contribution was significantly less to the CD spectra when the same concentration of protein solution was incubated with CBBG (Figure 2C). The bottom panel of figure 2B shows that the conversion of HI to β-sheet rich secondary structure was started at an early time and, subsequently it rapidly converted to β-sheet structure. Figure 2C shows CD spectra taken at different time interval of HI (320 μM) incubated with much lower concentration (30 μM) of CBBG. It shows that the α-helical pattern of the protein was preserved for a longer time (lower panel of Figure 2C) compared to protein alone (lower panel in Figure 2B). All these observations indicated a significant stabilization effect powered by CBBG and it can preserve the α-helical folds of HI for a long time.

**Figure 2.**
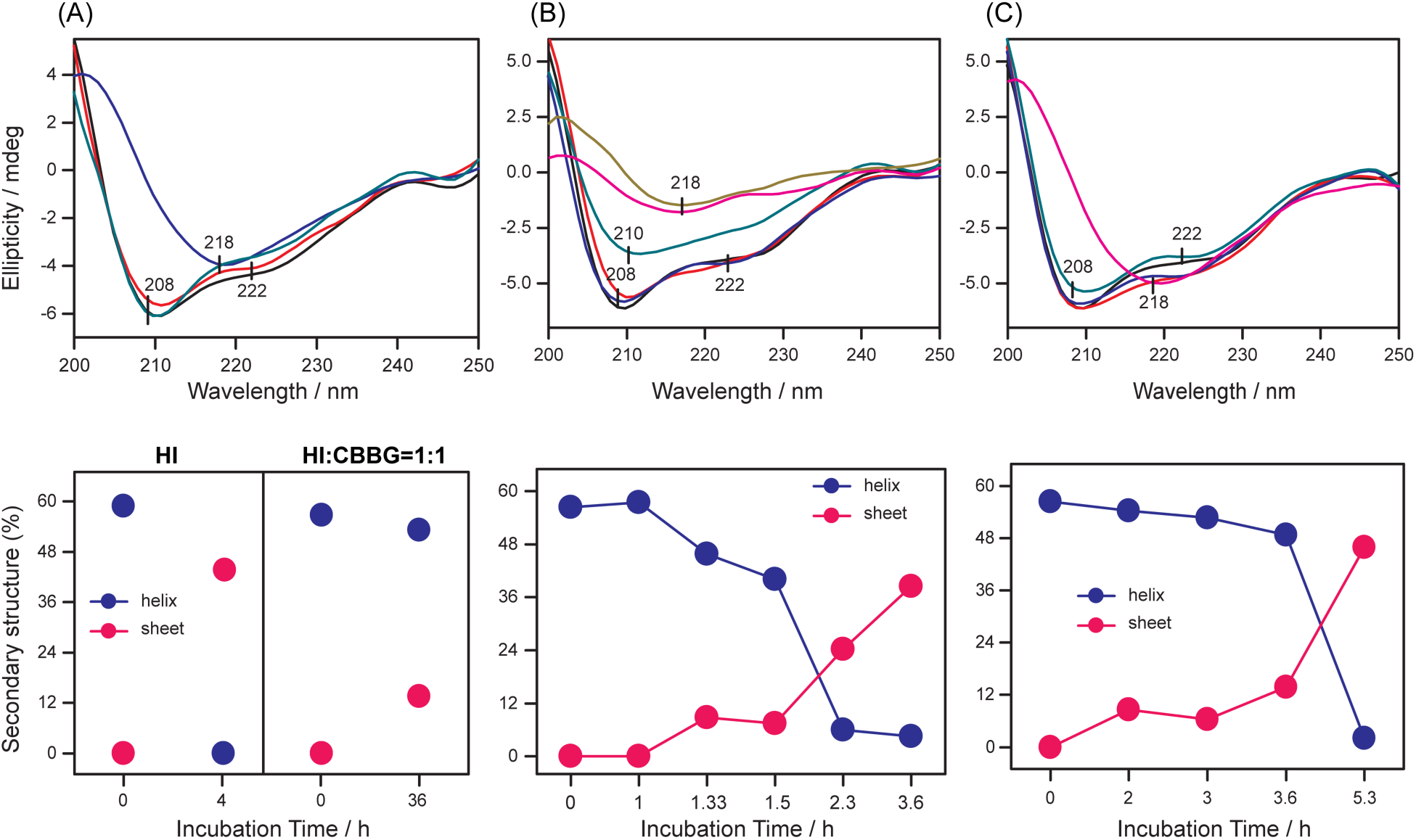
(A) Upper panel shows the CD spectra of HI (100 μM) in the absence and presence of CBBG (100 μM) at different time point of incubation (T = 60 °C, 25 mM HCl, 100 mM NaCl, pH ~1.6); HI (black trace, 0 h), HI+CBBG (red trace, 0 h), HI (blue trace, 4 h), HI+CBBG (green trace, 36 h). Lower panel shows the changes in the secondary structural component (%) against incubation time: α-helix (blue) and β-sheet (red). (B) CD spectra of insulin solution at different time periods of incubation at high concentration (320 μM): black trace (0 h), red trace (1 h), blue trace (1.33 h), green trace (1.5 h), pink trace (2.33 h) and yellow trace (3.6 h). Lower panel shows the changes in the secondary structural component (%) against incubation time: helix (blue) and sheet (pink). (C) CD spectra of aggregating insulin (320 μM) in the presence of CBBG (30 μM) of selected time points: black trace (0 h), red trace (2 h), blue trace (3 h), green trace (3.6 h) and pink trace (5.3 h). Lower panel shows the changes in the secondary structural component (%) against incubation time: helix (blue) and sheet (red).

### The insulin-CBBG binding interaction is an enthalpy-driven process

The binding interaction of CBBG with HI was further studied by isothermal titration calorimetry (ITC) experiment. It indeed provides a direct measurement of thermodynamic parameters (enthalpy, entropy, binding constant, and stoichiometry) describing ligand binding to a macromolecule. Figure 3A shows the raw calorimetric data profile of interaction between CBBG and HI at 25°C. The upper panel shows the thermogram and each of the negative peak represents an exothermic process inferring that heat was released upon injection of the HI into CBBG solution as a function of time. The lower panel in the Figure 3A shows the plot of the integrated heat response obtained from the raw data plotted against the total volume of protein solution added to CBBG solution. The exothermicity of the calorimetry peaks as shown in both the panels in Figure 3A suggests a significant interaction between CBBG and HI and thus the binding interactions between CBBG and HI were mainly exothermic in nature. The binding isotherm (solid line in the lower panel in Figure 3A) of the HI-CBBG interaction was generated by fitting the thermal data points and obtained thermodynamic parameters are given in Table 1. CBBG binds to HI with a moderate affinity. The dissociation constant (Kd) was 12.5 μM and, Gibb’s free energy of binding (ΔG) was –11.1 kcal/mol. CBBG formed a 1:2 complex with HI as judged from the fitting parameter value of ‘n’ (see Materials and Method) and the changes in enthalpy (ΔH) and entropy contribution (TΔS) were −57.2 and −46.1. kcal/mol, respectively. Thus, the ΔH and ΔS both were negative, and the binding was entropically disfavoured and the association of CBBG with HI was driven mainly by enthalpy contributions. The negative value of ΔH and ΔS further suggested that the binding of HI to CBBG may be dominated by hydrogen bonding and van der Waals interaction. NMR and molecular docking analysis as discussed in the subsequent sections, indeed, supported this finding.

**Table 1:** Binding constants and thermodynamic of binding between CBBG and insulin form ITC and NMR experiment.

| Complex | n | Method | Kd ($\mu\text{M}$) | $\Delta G(\text{kcal mol}^{-1})$ | $\Delta H(\text{kcal mol}^{-1})$ | $-T\Delta S (\text{kcal mol}^{-1} \text{K})$ |
| --- | --- | --- | --- | --- | --- | --- |
| Insulin-CBBG | 2 | ITC | 12.5 | -11.1 | -57.2 | 46.1 |
| | | NMR | $7.50 \pm 1.56$ (aliphatic region) | | | |
| | | | $5.26 \pm 1.64$ (aromatic region) | | | |
| | | | $7.54 \pm 2.56$ (amide region) | | | |

**Figure 3.**
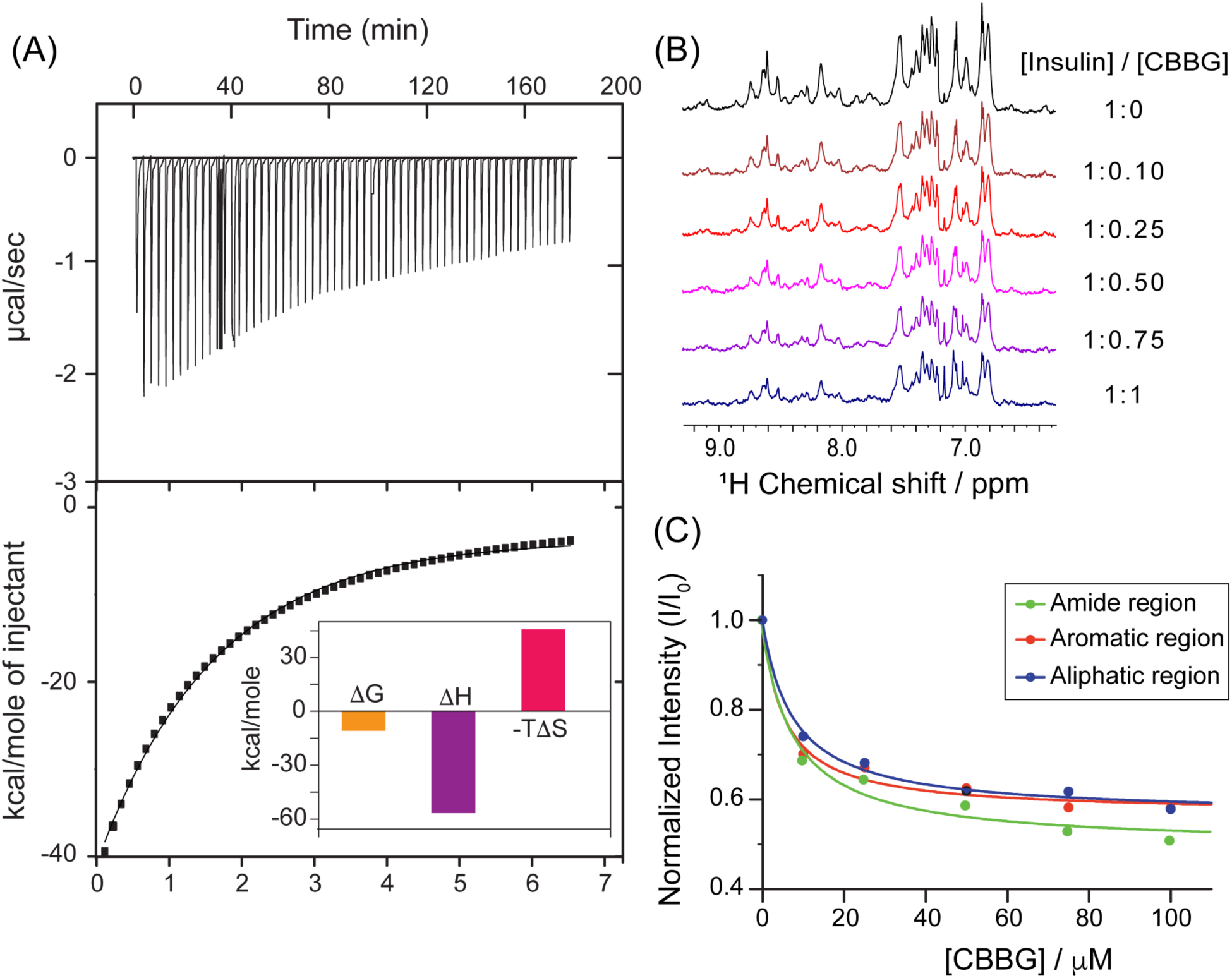
(A) ITC thermogram profile for insulin binding to CBBG. The upper panel presents the raw heat effects against time for the titration of CBBG (0.01 mM) with insulin (0.4 mM) in aqueous HCl solution (25 mM HCl, 100 mM NaCl, pH ~1.6) at 25 °C. The bottom panel shows the integrated heat data after correction of the heat of dilution against the molar ratio of CBBG to insulin. The black line indicates the fitted curve assuming a one-site binding model with one type of site. The binding free energy (Δ*G*), enthalpy (Δ*H*), and entropy (Δ*S*) of insulin to CBBG is shown in inset. (B) ^1^H NMR spectra of 100 μM insulin in 25 mM HCl buffer with 100 mM NaCl (pH ~1.6, black curve) and the presence of a different concentration of CBBG at 25° C. The concentration of CBBG were varied from 10 μM to 100 μM. (C) The NMR proton peak intensity decay of aliphatic (blue) aromatic (red) and amide (green) region of insulin upon titration with CBBG.

We performed a series of 1D NMR experiments for additional support and understanding the stabilizing effect of CBBG on HI. Figure 3B displays the 1D ^1^H NMR spectrum of HI (100 μM HI in 25 mM HCl buffer (pH ~1.6) containing 100 mM NaCl) in the presence of different concentration (0, 10, 25, 50, 75 and 100 μM) of CBBG. The protein alone exhibited distinct sharp peaks in the amide, aromatic and aliphatic regions (Figure 3B). However, the addition of CBBG into the solution of HI resulted in concentration-dependent broadening and, reduction in overall peak signal intensity of many proton resonances (Figures 3B and 3C). The uniform loss of signal intensities upon increasing concentrations of CBBG suggested upon a fast to intermediate exchange between the free and CBBG-bound HI at the NMR time scale.^21,44^ The intensity decay rate constants (Kd) of amide, aromatic and aliphatic regions (upon CBBG binding) were determined as 7.54 ± 2.56 μM, 5.26 ± 1.64 μM and 7.50 ± 1.56 μM, respectively (Figure 3C, Table 1). The binding affinity determined from NMR was in close agreement with the ITC determined Kd values (12.5 μM) (Table 1). Parallel to this, we also observed the changes in relative NMR signal intensities of the free and bound CBBG peaks upon titration with HI (Supporting Figure S2). Addition of 10 μM of HI fibril to CBBG (500 μM) revealed nearly 16% and 11% reduction in the 1D NMR signal intensities of the aromatic and aliphatic regions of CBBG (Supporting Figures S2B and S2C). With the addition of 25 μM HI to CBBG at 1:20 molar ratio we observed almost negligible chemical shift perturbations but broadening in the highlighted region (Supporting Figure S2), indicating a change in the chemical environment of CBBG. We also attempted transferred NOESY (trNOESY) experiments. However, since CBBG is a rigid molecule, we could not get a sufficient number of trNOE peaks (data not shown).

### CBBG binding prevents the formation of ‘partial unfolded’ state required for nucleation of HI fibrillation

The above observations (ITC and 1D NMR results) clearly showed that CBBG can bind to HI with micromolar affinity and stabilized the helical folds (CD analysis) of the protein. We further examined whether the binding added enough structural stability that could hinder the ‘partial unfolding’ of the compact helical structures, as observed by several other studies.^10,27,43^ Two experimental methods were employed for the same: (i) thioflavin T (ThT) fluorescence assay measurement that can detect fibrillation of the protein and (ii) 2D NMR analysis. ThT is a small dye molecule that becomes highly emissive upon binding to the cradle of the cross-β-sheet present in amyloid fibrils and produce a typical fluorescence emission spectrum with a peak maximum at ~482 nm in aqueous solution.^45^ However, in the absence of amyloid fibrils, it fluoresces very weakly. The growth curve (kinetics) was made by measuring fluorescence intensity of ThT in the presence of a quantitative amount of HI samples at different time points of incubation.

Figure 4A displays the time evolution of HI fibrillation based on ThT fluorescence values after incubation of HI (100 μM) at 60 °C (pH 1.6, 25 mM HCl, 100 mM NaCl buffer) and it exhibited a typical sigmoid curve comprising of an initial lag phase, in which nothing apparently occurred and, very little amount of amyloid fibril was formed. The presence of baseline level fluorescence represents lag phase and the enhancement in ThT fluorescence suggest the initiation of growth phase and the saturation in ThT fluorescence indicate the completion of fibril formation. The lag phase duration and apparent rate constant (*Kapp*) of fibrillation were derived from Equation 3 (Materials and Methods section). The resulting time duration/lag phase was found to be as ~1.8 h and suggested that no major increment of compact β-sheet structure in the incubated samples for this period of time (Figure 4A, Table 2). Subsequently, the growth phase started, and finally, it reached to a saturated phase within 2 h and the apparent rate constant for HI fibrillation was calculated as 0.83 h^-1^ (Figure 4A, Table 2). The fibril formation was also confirmed through AFM images (Figure 5). However, prior to formation of matured fibril it produced oligomer morphology (incubation for 60 min, Figure 5A) and subsequently it produced protofibril structure (Figure 5B). Eventually, after 200 min of incubation, these protofibrils organized to form long, compact, and dense fibrils with ~10-12 nm diameters (Figure 5C). Thus, the aggregation kinetics of HI followed a typical sigmoidal nature containing a lag phase association with nucleation and suggested the nucleation-elongation mechanism of fibril formation.

**Table 2:** Lag time and Growth Rate Constants for Insulin amyloid fibrillation formation (different concentrations and conditions see in Method Material) and in the presence and absence of CBBG).

| condition | reaction | lag phase | apparent rate<br>($k_{app}$) |
| --- | --- | --- | --- |
| Insulin 100 $\mu\text{M}$, pH $\sim 1.6$ | HI | 1.8 h | 0.83 $\text{h}^{-1}$ |
| Insulin 320 $\mu\text{M}$, pH $\sim 1.6$ | HI | 1.6 h | 0.95 $\text{h}^{-1}$ |
| CBBG1 7.5 $\mu\text{M}$ | HI/CBBG1 | 3.06 h | 0.42 $\text{h}^{-1}$ |
| CBBG2 15 $\mu\text{M}$ | HI/CBBG2 | 3.32 h | 0.42 $\text{h}^{-1}$ |
| CBBG3 30 $\mu\text{M}$ | HI/CBBG3 | 3.68 h | 0.32 $\text{h}^{-1}$ |
| Insulin 100 $\mu\text{M}$, pH $\sim 7.2$ | HI | 1.6 day | 0.88 $\text{day}^{-1}$ |

**Figure 4.**
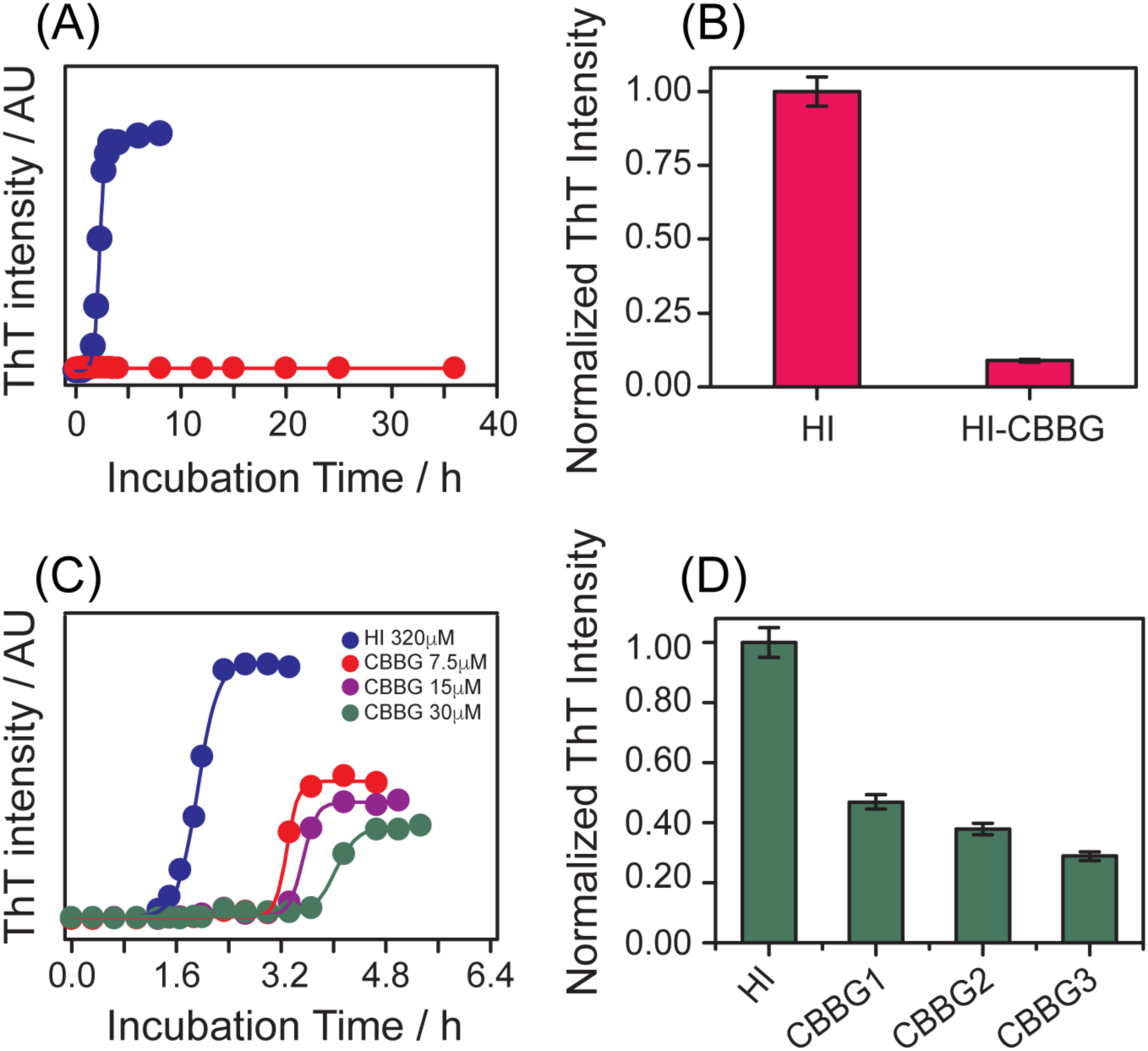
ThT fluorescence assay of insulin (HI) fibrillation in different solution conditions (details are provided in the Materials and Methods section): (A), ThT fluorescence intensity at 482 nm in the presence of the quantitative amount of incubated HI solution at a different time point of incubation. Incubation condition, 100 μM insulin, temperature 60 °C, 25 mM HCl, 0.1M NaCl, pH ~1.6, without CBBG (blue trace) and in the presence of 100 μM CBBG (red trace); (B), normalized (with respect to 0 concentration of CBBG) ThT fluorescence intensity (at 482 nm) at equilibrium (after five hours of incubation) of insulin incubated in the absence and presence of CBBG, other solution conditions are same as in A; (C), plot of ThT fluorescence intensity against time, similar to (A) for HI incubated at higher concentration (320 μM) and, in the absence and presence of a different amount of CBBG (7.5, 15 and 30 μM). (D) Normalized (with respect to 0 concentration of CBBG) ThT fluorescence intensity at equilibrium of the samples described in (C) and marked as HI (HI+ 0 μM of CBBG), CBBG1 (HI + 7.5 μM CBBG), CBBG2 (HI +15μM CBBG) and CBBG3 (HI +30 μM CBBG).

**Figure 5.**
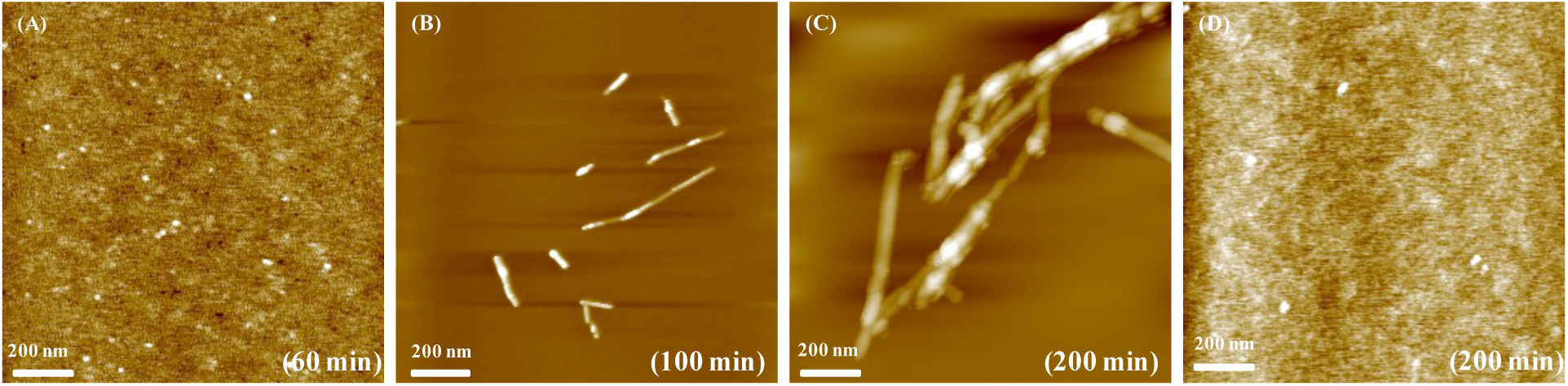
Representative AFM images showing aggregate morphology at several time point of HI (100 μM) incubation (temperature 60 °C, 25 mM HCl, 0.1M NaCl, pH ~1.6) in the absence and the presence of CBBG (100 μM). (A), (B) and (C) represent the surface morphology of the aggregates formed at 1 h, 1.6 h and 3.6 h, respectively, of insulin samples incubated without CBBG. (D) Represent the morphology of HI in the presence of CBBG at 3.6 h and no effective aggregates could be found. Scale bar is 200 nm.

However, HI (100 μM) solution co-incubated with CBBG (100 μM) at 1:1 molar ratio, growth phase would not appear (red curves in Figure 4A) and fibril formation was not detected Figure 4B shows very weak ThT fluorescence intensity of HI co-incubated with equimolar CBBG concentration. A relatively higher enhancement of ThT fluorescence intensity was observed after 3 h of incubation for HI incubated alone. Incubation in the presence of CBBG was continued for 36 h and no increase in the ThT fluorescence intensity was observed (Figure 4A). Also, under AFM, we could find no detectable oligomers or fibrillar aggregates from HI incubated in the presence of CBBG (Figure 5D). We further checked the attenuation effect of CBBG on HI fibrillation at ambient pH of ~ 7.2. Figure S3 shows the ThT fluorescence profile that represent the aggregation kinetics of HI solution incubated at pH of 7.2. It also showed a sigmoidal curve with three phases: lag phase, a succeeding growth phase followed by a plateau. In the absence of CBBG, the lag phase was found to be 1.6 days and the apparent rate constant of the fibrillation was 0.88 day^-1^ (Table 2). However, the ThT fluorescence enhancement was not very high and the AFM images confirmed that the aggregates formed at this pH was amorphous in nature (Figure S4). Figure 4C shows the kinetics of HI fibrillation at higher concentration (320 μM) (keeping other solution conditions same i.e. temperature 60 °C, buffer 25 mM HCl, 100 mM NaCl, pH ~1.6) in the absence and in the presence of much low concentrations of CBBG (7.5 μM, 15 μM, 30 μM).

Table 2 listed the lag phase durations and the rate constants of the fibrillation. The ThT fluorescence intensity plot indicated that both the nucleation phase (lag phase) and the growth phase were affected; however, the concentration of the protein was ~40 times higher than the CBBG. The presence of excess concentration of protein compared to CBBG could allow the free protein molecules to aggregate in its way along the inhibition effect CBBG in the nucleation phase. Figure 4D shows that the relative amount of fibril obtained even in the equilibrium state of fibrillation in the presence of different amount of CBBG at much lower concentration was very low, while the protein concentration was significantly high. It was further noted that the fibril formation was not evidenced in 100 μM protein sample incubated under 5 μM CBBG solution (data not shown). A similar observation of no fibrillation was also observed for very low concentration (1 μM) of HI incubated with the same concentration of CBBG (data not shown). These observations suggested that CBBG was effectively interfering with the thermal unfolding and fibrillation of HI.

Further, to evaluate the effect of CBBG on understanding the molecular intricacy of the inhibition process, two-dimensional ^1^H-^1^H NOESY experiments were performed with HI in the absence and in the presence of CBBG molecule. Figure 6 displays the homonuclear 2D-NOESYspectra of 1 mM HI in 10 mM sodium phosphate buffer containing 10 mM NaCl at pH 2 before and after incubation at 60 °C for 24 h both in the presence and absence of equimolar CBBG molecules. Figure 6A shows a large numbeof HI NOE peaks (at room temperature, before heating) originating from the cross-relaxation between two protons within 5 Å of each other.^46^ The recorded spectrum was similar to previously published NOESY data recorded in the similar experimental conditions.^47^ Surprisingly, after 24 h of incubation of HI sample at 60 °C, all the NOE cross-peaks were completely broadened (Figure 6C), suggesting the formation of a larger assembly, which were beyond the detection in NMR time scale due to slower tumbling and rapid T_2_ relaxation.^21,47–50^ HI sample incubated with equimolar concentrations of CBBG at room temperature also showed NOESY spectra with numerous NOE peaks (Figure 6B). However, incubating the solution for 24 h at high temperature (60 °C) the NOESY spectrum (Figure 6D) was very similar to the spectra (Figure 6B) obtained before heating the solution. Additionally, 1D NMR time kinetics (Supporting Figure S5) showed 91% and 93% decay in signal intensity at aromatic and aliphatic regions for HI alone after 24 h incubation, while in the presence of equimolar CBBG the decay of NMR signal intensity were only 14% and 21%, respectively, suggesting the stabilization of HI in presence of CBBG (Supporting Figure S5C and S5D). Collectively, the data revealed that CBBG inhibits the formation of NMR-invisible assemblies affecting the overall aggregation kinetics.

**Figure 6.**
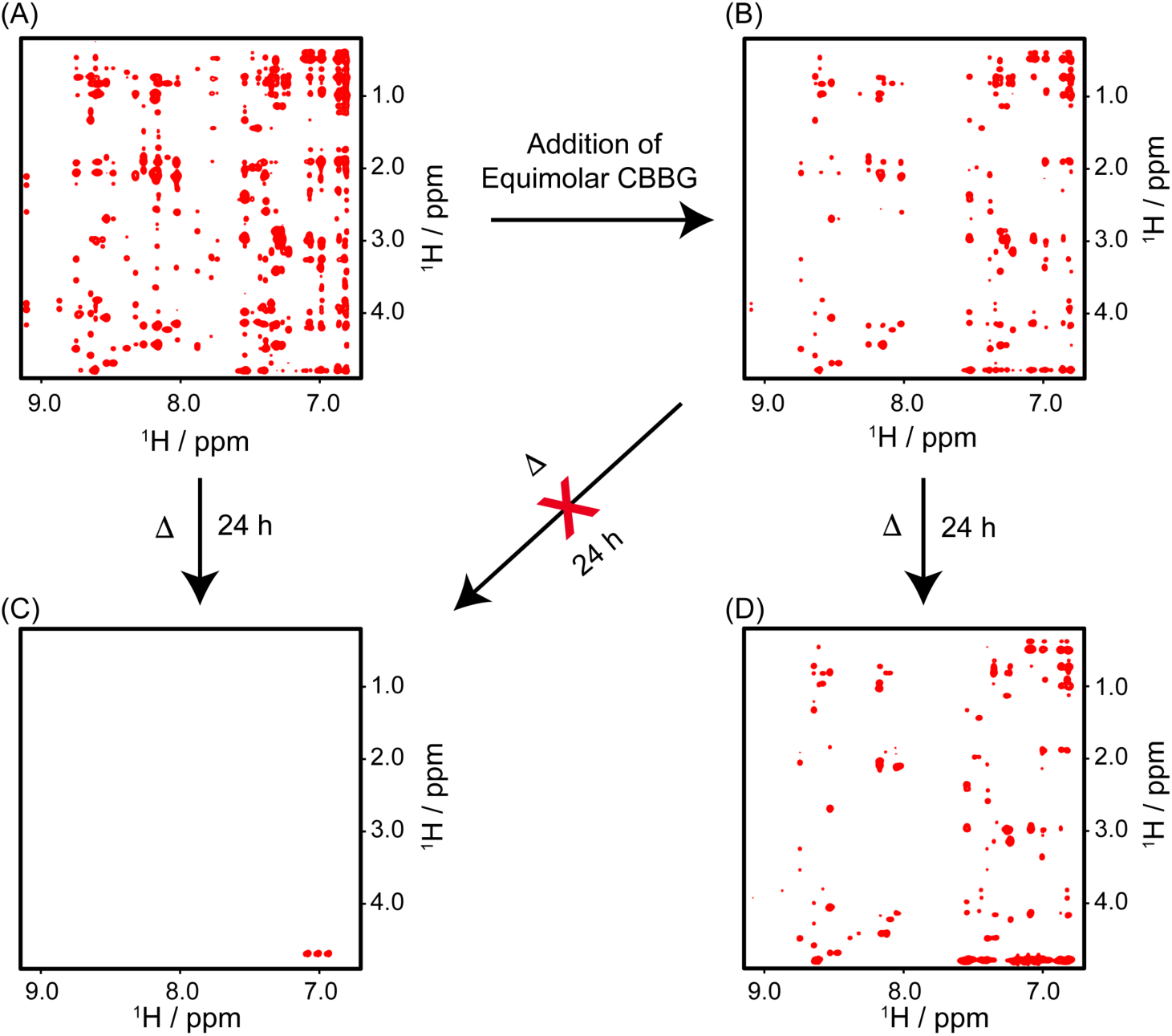
Inhibition of insulin amyloid fibrillation by CBBG. Two-dimensional NOESY NMR spectra: (A), insulin in the absence of CBBG and (B) in the presence of CBBG (B) just before incubation; (C) insulin solution after heating at 60 °C for 24 h shows no cross peak and (D) insulin co-incubated with equimolar CBBG at 60 °C for 24 h shows almost identical cross peak recorded before incubation (B). The experiment was performed in 10 mM sodium phosphate, 10 mM NaCl (pH=2.0) and 10% D_2_O using Bruker Avance III 700 MHz at 25°C.

### Binding epitope of CBBG on insulin that attenuates aggregation

HI remains quite stable in its zinc-bound hexameric structure. However, in the absence of zinc ion at physiological pH (~7.0) it is primarily present as a dimer and aggregates very easily at elevated temperatures. At pH ~ 2.0 and room temperature, HI forms dimmers (due to charge repulsion between Zn^2+^ and protonated His^B10^)^51^, as confirmed by solution-state NMR^43,47,52^ and X-ray crystallography.^9^ Dimeric structure of human insulin in sodium phosphate buffer (pH 2.0) was indicated by a series of intermolecular NOEs between the two monomers of the dimer (Supporting Figure S6). The inter-monomer NOEs of the dimer have been observed for Gly^B8^C_α_H/Tyr^B16^C_ε_H, Ser^B9^C_α_H/Tyr^B16^C_ε_H, Ser^B9^C_β_H/Tyr^B16^C_ε_H, Ser^B9^C_α_H/Tyr^B16^C_δ_H, Val^B12^C_γ_H/Tyr^B16^C_ε_H, Tyr^B16^C_β_H/Tyr^B26^C_δ_H, Gly^B23^C_α_H/Tyr^B26^C_δ_H, and Pro^B28^C_γ_H/Tyr^B16^C_ε_H (Supporting Figure S6). These NOEs corroborated well with the previous study.^47,52^ Apart from these, no other intermolecular NOEs between HI and CBBG were observed.

Figures 7A–B and S7 show the overlap of 2D NOESY NMR spectra for HI and HI/CBBG (1:1) complex at low pH (i.e., pH 2) before (Figure 7A and S7A) and after heating at 60 °C for 24 h (Figure 7B and S7B). Close inspection suggests that CBBG interaction causes several chemical shift perturbations (CSPs) as well as line broadening effects for the HI NMR peaks. A detailed analysis of the room temperature 2D NOESY spectra reveals that the several residues from both the HI chains (A and B) interact strongly with CBBG. Among the most perturbed residues in the A chain were Val^A3^, Glu^A4^, Cys^A7^, Thr^A8^ from the N-terminal α-helix and Leu^A16^ from C-terminal helix (Figures 7C and 7E). Similarly, Val^B2^, Asn^B3^, Ser^B9^ from the N-terminal region of B chain β-turn, Arg^B22^ from the second β-turn region and Phe^B24^, Phe^B25^ from C-terminal β-strand, confirmed by chemical shift changes in 2D NOESY spectra (Figures 7C and 7E). This indicates that HI undergoes a local (residue level) conformational change upon binding to CBBG.

**Figure 7.**
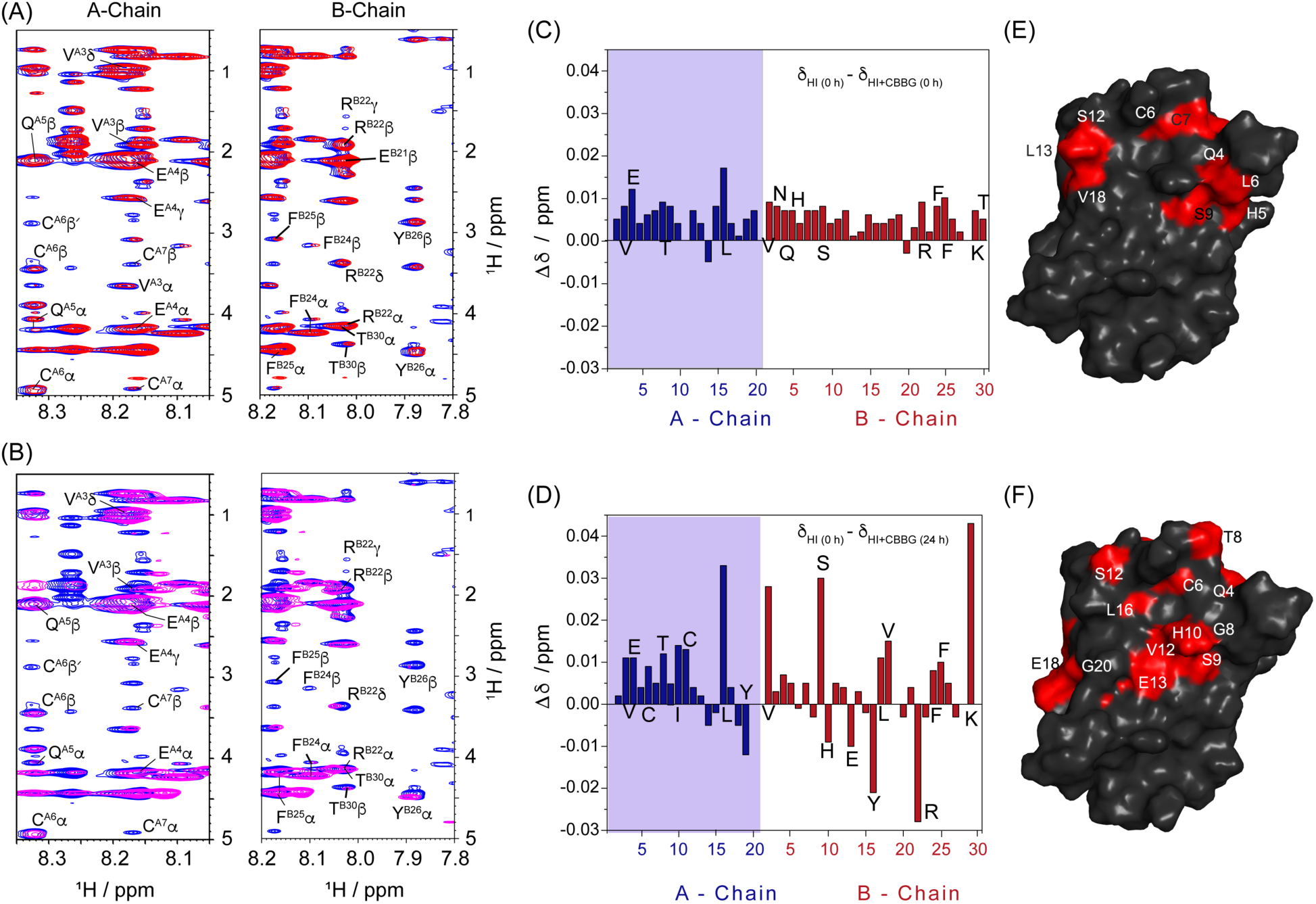
Structural insights into the insulin amyloid inhibitory interaction with CBBG. Two-dimensional NOESY NMR fingerprint region of interaction site of CBBG in insulin A and B-chains before (A) and after (B) incubation for 24 h at 60 ºC in 10 mM sodium phosphate (pH 2.0) containing 10 mM NaCl and 10% D_2_O. Blue, red and purple colours represent insulin (A and B), insulin-treated with equimolar CBBG (insulin:CBBG=1:1) before heating (A) and after heating (B) at 60 ºC, respectively. The changes in proton chemical shift (Δδ) of insulin residues were visible both in A-chain and B-chain of insulin, before (C) and after (D) heating of insulin complex (insulin: CBBG = 1:1) at 60 ºC for 24 h. Chemical shift change (Δδ) of alpha protons (α-H) were calculated from the difference between the chemical shift of respective residues of insulin (δ_HI_) at 0 h and insulin+CBBG (δ_HI+CBBG_) at 0 or 24 h, respectively. 2D-NOESY NMR spectra were recorded on Bruker Avance III 700 MHz, equipped with RT-probe. (E) and (F) are the mapping of chemical shift perturbed residues onto the surface of insulin dimeric structure at 25 °C and 60° C, respectively; clearly delineates and points to the binding cavity at the interface of A-and B-chains of insulin (at which CBBG could be docked).

Further, before longer incubation time with equimolar concentrations of CBBG (1:1 molar ratio) showed a line-broadening effect for Glu^A4^, Gln^A5^, Cys^A7^ and Tyr^A19^of the A-chain and Gln^B4^, Gly^B8^, Ser^B9^, Val^B12^, Val^B18^, Cys^B19^, Glu^B21^, Tyr^B26^ and Thr^B30^ from the B chain (Figures 7A and S7A). A parallel 1D NMR analysis of the spectra showed the percentage of signal intensity broadening (Supporting Figure S5B). Interestingly, the total proton signal intensity of HI was reduced to 0.71 unit (71%) in presence of equimolar CBBG concentrations (Supporting Figure S5B). The chemical shift perturbation and broadening of HI NMR peaks indicated the significant binding followed by stabilization of HI sample by CBBG.

Furthermore, the residue-specific interactions with HI dimer were also identified at aggregation-prone conditions i.e., by increasing the sample condition to a temperature of 60 °C (Figures 7B, S5 and S7B). The proton NMR spectra of HI upon incubation at high temperatures in the presence of CBBG showed more chemical shift perturbations, line broadenings and the appearance of new signals as shown in the highlighted region in Figure S5A. Interestingly, after 24 h of incubation at 60 °C the 2D-NOESY NMR spectra showed significant chemical shift changes for the Val^A3^, Glu^A4^, Cys^A7^, Thr^A8^, Ile^A10^, Cys^A11^, Leu^A16^ and Tyr^A19^residues from the A-chain and Ser^B9^, His^B10^, Glu^B13^, Tyr^B16^, Leu^B17^, Val^B18^, Arg^B22^, Phe^B24^, Phe^B25^ and Lys^B29^ residues from the B-chain of HI. This data, as represented in Figures 7D and 7F, suggests that the HI dimer undergoes a conformational change upon binding to CBBG at 60°C. Additionally, remarkable line broadening in presence of CBBG was observed for the Glu^A4^, Gln^A5^, Cys^A7^, Leu^A16^, Glu^A17^and Tyr^A19^ residues from HI A chain as well as Gln^B4^, Gly^B8^, Ser^B9^, Val^B12^, Glu^B13^, Val^B18^, Cys^B19^, Glu^B21^, Arg^B22^, Phe^B25^, Tyr^B26^, Lys^B29^ and Thr^B30^ residues from the B-chain (Supplementary Figures S7B and S8). These NMR data provided atomic resolution information for the specific location on the HI molecule where the interaction with CBBG takes place.

### CBBG Docking and pharmacophore mapping

To gain further insights into the mechanisms of these molecular interactions, we performed molecular docking of CBBG on HI based on the chemical shift perturbation (CSP) data of ambiguous interaction sites. Our results suggested that the CBBG was docked onto the cavity lined by the above highlighted residues at the interface of A and B chains of HI in one monomer with a very good PLP fitness of 82.5 (Figure 8A and 8B). Chemically, CBBG bears a N, N-disubstituted tri-anilinyl-methane scaffold and with an IUPAC chemical name as 3-(((4-((4-((4-ethoxyphenyl)amino)phenyl)(4-(ethyl(3- sulfobenzyl)amino)-2-methylphenyl)methyl)-3-methylphenyl)(ethyl)amino)methyl)benzene sulfonate] (Figure 1C). Among the three aniline fragments on CBBG, one fragment - N-ethoxy-phen-4-yl-aniline, wherein both the aromatic phenyl rings and the N-ethoxy moieties in this fragment were engaged in hydrophobic interaction with Ile^A10^ (5.46 Å, 4.63 Å, respectively, Supplementary Figure S9) residues of insulin A subunit. Interestingly, the other two fragments N-ethyl-N-(phenyl-3-sulfonic acid)-2-methyl-aniline were involved in strong hydrophobic interactions with A and B subunits, respectively (Figure 9). One of the hydroxyl group of sulfonic acid moiety on the N-phenyl group oriented towards insulin A subunit was involved in hydrogen bonding interaction with main-chain carbonyl atoms of N3 (2.30 Å). While the ‘-CH’ atoms on N-methyl-phenyl moieties were mediating close C-H contacts with carbonyl atoms of main chain Cys^A6^ (2.64 Å) and strong alkyl and aromatic hydrophobic interactions with Cys^A7^ (3.86 Å, 4.95, respectively). Further, 2-methyl-group on aniline fragment was involved in strong alkyl hydrophobic interaction with Cys^A6^ (4.49 Å), Cys^A11^ (4.46 Å), Leu^A16^ residues. The aromatic ring of aniline on this fragment was mediating alkyl hydrophobic interaction with Leu^B11^ (4.87 Å, Supplementary Figure S9).

**Figure 8.**
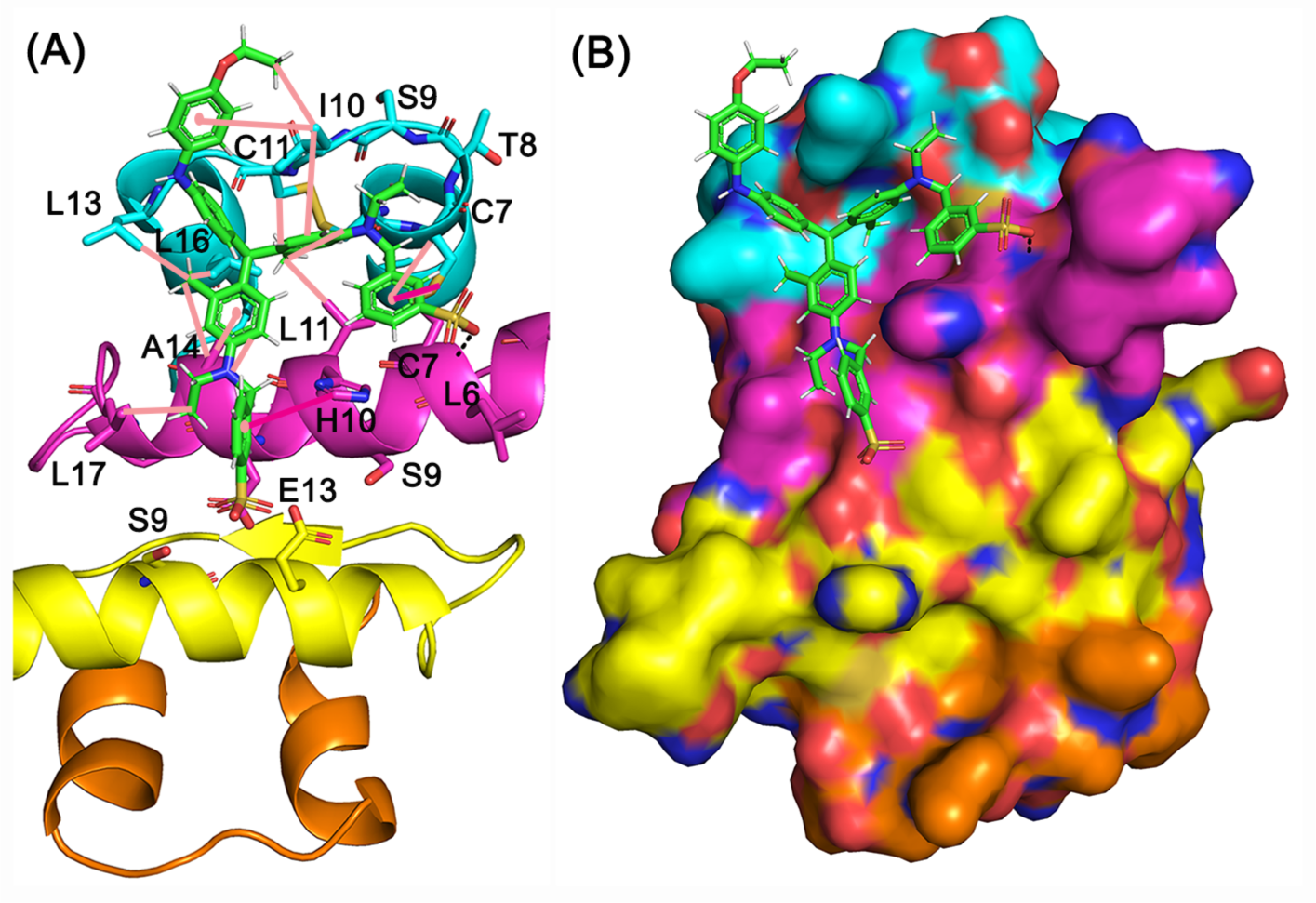
Binding model of CBBG at the dimeric interface of insulin A, B subunits. (A) Cartoon model of CBBG binding highlighting the predominant network of alkyl hydrophobic interactions (pink lines) and aromatic interactions (magenta). (B) Surface view of CBBG best pose at the dimer interface.

**Figure 9.**
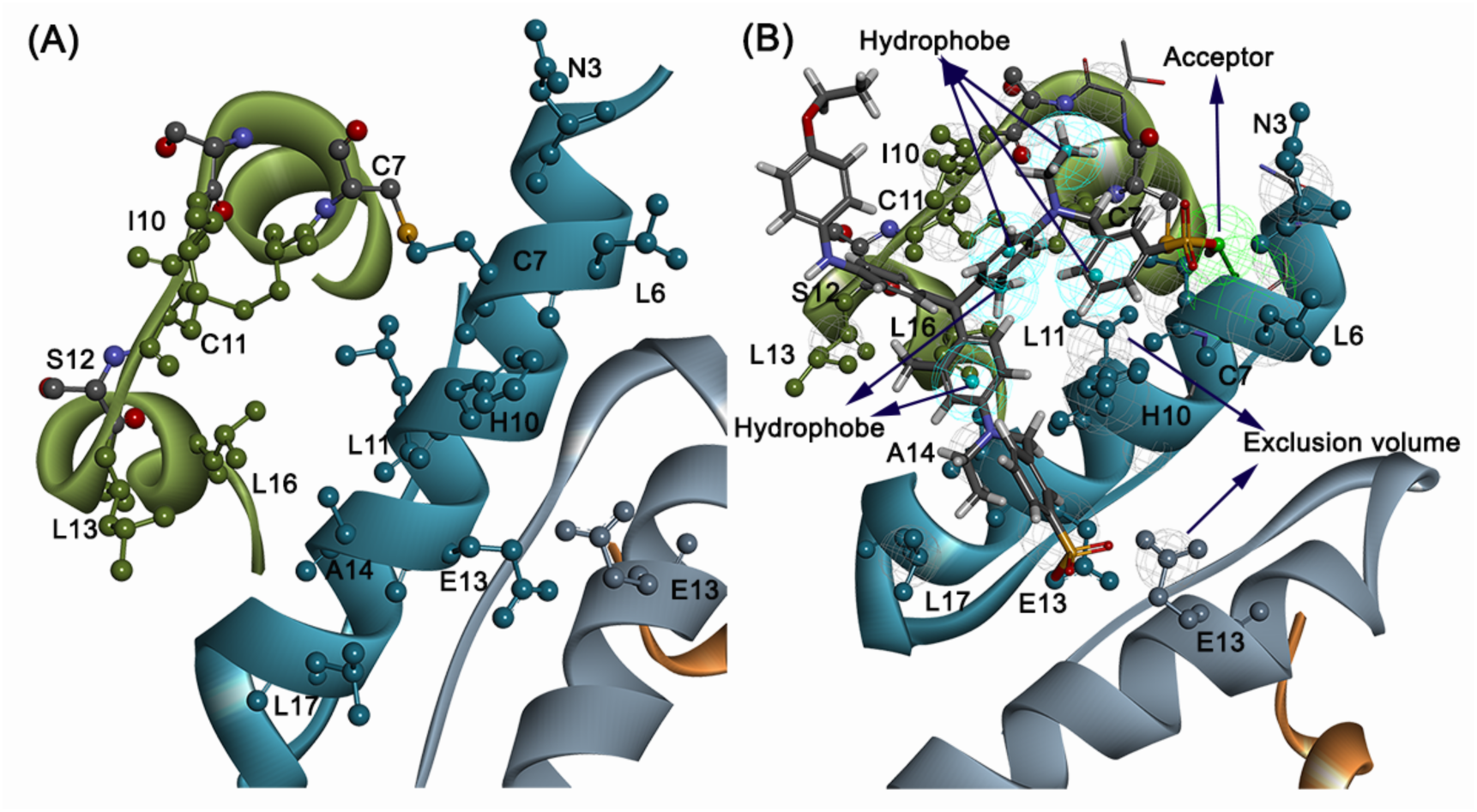
Mapping of important residues involved in CBBG interaction and its pharmacophore model. A) Residues highlighted from NMR and Molecular docking interactions. B) Receptor ligand-based pharmacophore model reveals the five hydrophobic features at the AB interface and acceptor feature in the vicinity of C7 and N3 residues.

Similar deep network of both aromatic and alkyl-aromatic was seen predominantly with the other N-ethyl-N-(phenyl 3-sulfonic acid)-2-methyl-aniline fragment positioned into the B subunit of HI. The terminal N-ethyl, N-N-(phenyl-3-sulfonic acid)-moieties on 2-methyl-aniline were engaged in alkyl hydrophobic interaction with Leu^B17^ (4.62 Å) and aromatic interaction of (phenyl-3-sulfonic acid)- with His^B10^ (5.06 Å). The sulfonic acid was not positioned favourably to mediate H-bonding interaction, though it was in close vicinity with Glu^B13^ from both B subunits in the dimer (4.89 Å, 3.38 Å respectively, Figure 9). Lastly, the 2-methyl-aniline ring was involved in alkyl hydrophobic interaction with Leu^A13^ (5.02 Å), Leu^A16^ (5.46 Å) and the phenyl ring of aniline with Ala^B14^ (3.50 Å, Supplementary Figure S9). Taken together, the strong network of hydrophobic interactions, close contacts and hydrogen bonding interaction at the interface of A and B subunits of HI stabilize the ligand binding and enhance the stability from aggregation.

### Pharmacophore Mapping

Our NMR results and molecular docking interaction studies have enabled us to map the pharmacophore features at the very important residues involved in interaction with CBBG. Thus, six receptor-ligand based pharmacophore features comprising of five hydrophobic features anchoring Cys^A7^, Ile^A10^, Cys^A11^, Leu^A16^ residues and Cys^B7^, Leu^B11^, His^B10^, Ala^B14^ residues and an acceptor feature in the vicinity of Cys^B7^ and Asn^B3^ residues of HI were mapped (Figure 9). The exclusion volumes encode the broad contour of protein active site shape and help to prevent clashes of ligand atoms with protein atom. In future works, this vital pharmacophore model on HI protein does help us in identifying novel chemical entities that could prevent HI aggregation and stabilize HI shelf life.

### Can CBBG interact with monomeric human insulin?

It is reported that HI exists as a monomer in 20% acetic acid.^21,43,44^ To identify the binding epitopes of monomeric HI by CBBG (HI:CBBG=1:1), we, again, recorded the NMR spectra of HI in 20% acetic acid-d_4_ (pH=1.9) at room temperature (Supplementary Figures S10 and S11). 1D and 2D NOESY NMR spectra of HI in the presence of equimolar CBBG revealed almost no significant chemical shift changes and line broadening effect, indicating that addition of CBBG did not induce any structural conformational changes within the residues of HI (Supplementary Figures S10 and S11). A few residues namely Cys^A11^ and Ser^A12^ (C-terminal helix region) from A-chain and Leu^B15^ and Phe^B25^ from B-chain display negligible chemical shift changes (<0.007ppm) upon equimolar CBBG addition (Supplementary Figure S11). Collectively, these results indicated that CBBG did not prefer to bind significantly to HI monomer. We further aimed to determine the binding epitope of HI at neutral pH. However, weak solubility of HI in phosphate buffer (pH ~7) hindered to perform similar NMR experiments in neutral pH. All the above-mentioned NMR experiments confirmed that the CBBG molecule can interact with dimeric HI and stabilizes the helical structure of HI

## Discussion

Protein amyloidogenesis has been a pathophysiological phenomenon underlying several of the known degenerative disorders, including Alzheimer’s disease, Parkinson’s disease, Type 2 diabetes, etc.^53^ Despite the advances, the exact mechanism of aggregation invivo, remains far from being elusive, majorly owing to the complexities and heterogenous properties of the different protein systems. Decades of amyloid research have been dedicated to delineating the intermediate oligomeric or proto-fibrillar stages that form the pathogenic conformers underlying disease etiology. However, these intermediates form a dynamic heterogeneous pool of several transient conformers, significantly different from the well-defined fibrillar forms, and are difficult to characterize or delineate.^23,54^ Without the atomic-level information on the growth of amyloid fibers, it has been challenging to design inhibitors or excipients to block fiber growth. In this context, it is indispensable to discuss the far-reaching possibilities of NMR experiments to enable the indirect probing of the nucleation events in the amyloidogenic pathways.^33,55^

The protein-folding intermediates have been implicated in harbouring the essential chemical cues that provide useful insight into understanding the amyloidogenic propensity. ^33,55^ These intermediates are often structurally similar to the low molecular weight early conformers of the protein but form a chemically distinct pool that nucleates the early aggregation events. High-resolution NMR spectroscopy has helped us to unveil the mechanism of Aβ amyloidogenesis in Alzheimer’s disease to be actually a multi-step process involving the docking of the free monomeric forms to the nucleating intermediates.^55,56^ Similarly, in Parkinson’s disease, atomic-resolution characterization of the α-synuclein protein enabled us to probe the “open” conformation of the monomeric form that serves as the intermediate in initiating the amyloidogenic cascade.^56^ Previous studies have also indicated that the amyloid fibril formation for globular proteins, such as HI, occurs via partially unfolded intermediates that gradually associate to form the oligomers, eventually into well-ordered mature fibrils.^9,14,23,27,33,47,57,58^ A partial unfolding of the monomeric globular protein produces a molten globule like structure that is comparable to the amyloidogenic protein intermediates, acting as the nucleating form.^29,50,55,59–61^

In the absence of zinc at physiological pH, HI is primarily present as a conformationally stable dimer; however, it can aggregate at elevated temperatures or during prolonged storage. Coarsely identical to the initial monomeric structure of the protein in buffer solution, these monomeric structures produce the lesser known nucleus *en route* to HI fibrillation.^9,14,23,27,33,34,47,57^ The nucleation step has been identified with the C-terminal end of the B chain helix turning away from the C-terminal end of the A chain helix. This causes a partial loss of helicity on the C-terminal of the A-chain, accompanied by a compromise of the intra-chain hydrophobic contacts between Leu^16A^ on A chain C-terminal helix and Leu^15B^ and Val^18B^ on the B chain helix. The consequences of the loss of these contacts is an opening up of the structure that creates a hydrophobic cavity in the centre of the protein comprised of the Phe^24B^, Phe^25B^, Tyr^26B^, Val^12B^, Val^3A^, and Tyr^19A^ residues that may serve as a nucleation site for the very early step of fibrillation.^57^ The NMR analysis defined several residues in the α-helical folds of the A chain along with other residues from either terminal segments of both the A and B chains that partake in producing the hydrophobic cavity, prompting the oligomerization events. This atomic-resolution characterization has been imperative in enabling us to define efficient pharmaceutical excipients to arrest the aggregation-pro conformer.

The present study suggested the role of CBBG, a close analog of brilliant blue FCF, used as an FDA approved food colour, in a stable binding to the epitope of molten globular intermediate formation in HI aggregation. CBBG induced perturbations of the pivotal unfolding events effectively arrest HI in the low molecular weight conformations. Interestingly, CBBG addition at 60 °C also showed close proximity (NOE contacts) to the Glu^13B^ of the HI B chain, which has been suggested to partake in the dimerization of the domain.^57^ Insulin B chain has been shown to adopt a β-hairpin topology wherein the N and C-terminal ends are held together by hydrophobic contacts and form a homodimer being held together by Glu^13B^. This dimeric β-hairpin topology allows the B chain to interact with A chain residues in the native folds of HI. Glu^13B^ residues of HI come to close contact with CBBG because of structural rearrangement during high temperature long incubation at pH 2. The close proximity of sulphonic acid group of CBBG and Glu^13B^ residues, suggests a strong interaction between them, as evidenced from the perturbed chemical environment of Glu^13B^ residues in the CSP data and line broadening of Glu^13B^ residues. This interaction prevents the unwinding of CTD and stabilizes the HI dimer structure. The NMR-based atomic resolution data collected, and molecular docking studies suggested that the CBBG binds to HI at the dimeric stage and prevents the availability of the crucial residue segments that partake directly in further oligomerization.

Our results showed that CBBG establishes a strong network of hydrophobic interactions, close contacts, and hydrogen bonding interactions between the two HI chains. These interactions were used to map the pharmacophore features of this intricate CBBG interacting interface that enhance the conformational stability of the arrested intermediates. We strongly believe that the proposed six feature receptor-ligand based pharmacophore model would help in identifying novel chemical entities to be incorporated as part of HI formulations to increase its shelf-life and pharmacological applications.

## Conclusion and Perspectives

Our studies demonstrate the rationale behind the usage of small molecules such as CBBG in the production, storage, and delivery of pharmaceutical formulations of human insulin in its most active form. The enthalpy driven binding of CBBG to HI with micromolar affinity allows stabilization of its α-helical form, preventing further unfolding of the protein that serves as the nucleation step in HI fibrillation. The residue-specific interactions provide a clear understanding of the stepwise mechanism of interaction. The ability of CBBG in binding to the specific domains of the dimer that prevent the native unfolding of the protein, hints at its immense potential as an excipiant in insulin formulations. NMR results and computational analysis further lead us to propose a ligand based pharmacophore model comprised of 5 hydrophobic and a hydrogen bond acceptor features that can anchor the residues at A and B chains of insulin. In future works, this vital pharmacophore model on HI protein does help us in identifying novel chemical entities that could prevent HI aggregation and stabilize insulin’s shelf life. Thus, our observations suggest upon a promising use of CBBG in the efficient production, storage and bio-availability of the active form of hormonal protein insulin.

## Materials and Methods

### Materials and sample preparation

Human HI (HI, 91077C), Thioflavin T (ThT), Coomassie brilliant blue G-250 (CBBG), NaCl and HCl were purchased from Sigma-Aldrich. Milli-Q water was used in the preparation of a buffer and stock solution of ThT dye. HI solution was prepared in 25 mM HCl containing 100 mM NaCl (pH ~1.6) to form a stock solution and then centrifuged at 15000 rpm for 10 min and passed through a 0.22 μm pore size filter to remove any insoluble aggregates. The stock concentration of the protein solution was determined through JASCO-600 UV-vis spectrophotometer, using the extinction coefficient value of the protein studied was as follow 6200 M^-1^ cm^-1^ at 276 nm. For inducing amyloid fibrillation, freshly prepared HI (320μM and 100 μM) solution in absence and presence of CBBG was incubated for several hours at 60 °C without agitation.

### Circular dichroism (CD) measurements

Far-UV circular dichroism (CD) spectra were recorded at 25 °C on a Jasco J-815 spectropolarimeter (Easton, MD). For CD analysis, freshly prepared HI (320 μM and 100 μM) solution in the absence and presence of CBBG was incubated for several hours at 60 °C without agitation. HI solutions were prepared in 25 mM HCl containing 100 mM NaCl (pH ~1.6). The spectra were measured using diluted aliquots (final concentration was ~ 15 μM of HI) at a different time point of incubation. 300 μl of the incubated protein solution was taken in 0.1 cm path length cuvette and scanned between 200–250 nm with a scanning speed 50 nm/min, resolution of 0.2 nm. For each sample, the representative spectrum was average of at least three individual scans.

### Isothermal Titration Calorimetry (ITC)

ITC measurements were carried out at 25 °C on a VP-ITC titration microcalorimeter (Micro Cal Inc., Northampton, MA). HI and CBBG samples were thoroughly degassed on a thermovac before the use in titration. For the correction of heat of dilution, the sample cell was loaded with buffer (pH ~1.6). and the reference cell was also filled with the same buffer solution. The solution in the cell was stirred at 90 rpm by the syringe filled with 0.4 mM HI in identical buffer solution. Injections of 4 μl of the buffer in the syringe were started after stability in baseline reached. 28 sequential such injections were made into the ITC cell containing buffer solution. The titration of the CBBG solution (0.01 mM) in the cell was followed by 28 sequential 4 μl injections of 0.4 mM HI into the ITC cell containing 1.8 ml of CBBG solution. The protein and CBBG solution were made in identical buffer condition (25 mM HCl containing 100 mM NaCl (pH ~1.6)). The raw calorimetric data profile (heat released) of interaction between CBBG and HI at 25°C were collected automatically and subsequently fitted to a one-site binding model by the Microcal LLC Origin 7.0 software. After subtracting the heat of dilution, a non-linear least-squares algorithm was used to fit an equilibrium binding equation to the data points (heat flow per injection against the concentration ratio of HI and CBBG). This best fit provides the apparent binding stoichiometry (*n*), the change in enthalpy (∆*H*), and the dissociation constant (Kd). The change in free energy (∆*G*) and change in entropy (∆*S*) for the binding reaction were analysed by the important equations of thermodynamics.

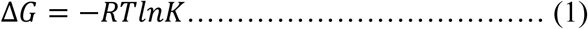

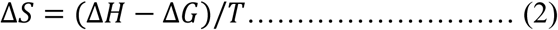

### ThioflavinT (ThT) Fluorescence Assay

ThT fluorescence is recorded to investigate the kinetics associated with fibril formation of human insulin (HI) in different solution conditions. The growth curve (kinetics) was made by measuring fluorescence intensity of ThT in the presence of a quantitative amount of HI samples taken at different time points of incubation of the HI solution in the presence and in the absence of CBBG. Buffer and other sample conditions were similar to samples used in ITC and CD measurements. 4 μl of incubated HI solution (either in the absence or presence with CBBG) was pipette out, added to 500 μl solution of ThT (~22 μM) and mixed carefully for the acquisition of fluorescence emission spectra using a Cary Eclipse fluorescence spectrophotometer. The optical path length of the fluorescence cuvette was 10 cm. The fluorescence emission wavelength range was 450-600 nm (excitation 440 nm, emission peak maximum ~ 482 nm). ThT fluorescence peak intensity at 482 nm was plotted against time, analysed and fitted to the sigmoidal curve using equation 3.^14^

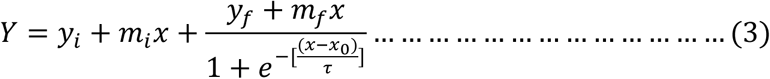

Where *Y* is the ThT fluorescence intensity at particular time (x), *x* is the incubation time, and *x*_0_ is the time to reach 50 % of maximal fluorescence; other parameters are determined by the fitting. The lag time is defined by *x*_0_− 2τ. Apparent rate constant (1/ τ), *m*_*i*_ and m_f_ are two constants (linear coefficients)

### Atomic Force Microscopy

Morphologies of the insulin aggregates produced in the absence and presence CBBG were performed using Pico plus 5500 ILM AFM system (Agilent Technologies). Incubation and sample conditions were similar to sample prepared for ITC and ThT fluorescence assay measurement described earlier. Micro-fabricated silicon cantilevers (resonance frequency of 300 kHz and spring constant range of 21-98 N/m) was used derive the morphological features of the aggregates formed from the incubated samples. The aliquot taken at defined time of incubation was diluted with water and drop-casting was made on freshly cleaved muscovite mica substrate. The solvent was removed by evaporation at room temperature in the open air. The images were captured with a scan speed of 0.5 lines/sec and processed using Pico view version 1.1 software (Agilent Technologies).

### ^1^H proton NMR

All NMR spectra were recorded on Bruker AVANCE III 700 MHz, equipped with RT probe and at 298 K. All NMR data acquisition and processing were done by using Topspin v4.0.6 software (Bruker). The interaction of CBBG dye with human HI (100 μM) in 25 mM HCl buffer with 100 mM NaCl and 10% D_2_O at pH ~1.6 was determined through a series of one-dimensional (1D) ^1^H proton NMR spectra of HI in presence of 0, 0.1, 0.25, 0.5, 0.75 and 1 molar excess of CBBG dye (from a stock of 3 mM CBBG). The NMR signal intensity data were fitted by using equation 4:

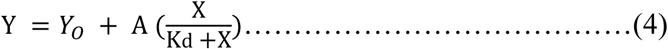

Here, Y is normalized signal intensity value, *Y*_*O*_ is initial signal intensity value, A is the fraction of fast exchange protons under our condition, X is the concentration of added CBBG dye (μM), and Kd the apparent intensity decay rate constant or fast exchange rate constant between free and bound CBBG.

Human HI (1 mM) was dissolved in 10 mM sodium phosphate buffer (pH 2.0) containing 10 mM NaCl and 10% D_2_O, with or without the addition of CBBG at 1:1 molar ratio for kinetics experiment using NMR. One-dimensional ^1^H and two-dimensional homonuclear ^1^H-^1^H NOESY NMR spectra (400 ms mixing time) were taken before and/or after incubation of samples at 60 °C for 24 h. At the same time, zinc-free human HI was prepared by the addition of EDTA followed by extensive dialysis and lyophilization. The lyophilized sample was then dissolved in 20% acetic acid-d_4_ and 10% D_2_O (pH 1.9). NOESY spectra of 350 μM HI were performed in the presence or absence of equimolar CBBG dye with a mixing time 200 ms at 25 °C. 512 increments in t_1_ and 2048 data points in t_2_ dimension along with excitation sculpting pulse sequence were used for water suppression for all NOESY spectra. The NOESY spectra of HI were also recorded at 25 °C with 64 scans and a spectral width of 12 ppm for both the dimension. The transferred NOESY (tr-NOESY) experiment (mixing times= 150 and 250 ms) of CBBG was recorded in the presence of HI fibril at a molar ratio of 1:20 (HI: CBBG, keeping all parameters constant. Parallelly, we also recorded 1D ^1^H NMR of 500 μM CBBG with the treatment of 10 μM and 25 μM HI fibrils in 25 mM HCl containing 100 mM NaCl and 10% D_2_O (pH ~1.6) at 25°C.

### Molecular docking

Dimeric assembly of insulin A and B subunits with PDB ID 2MO1^40^ was utilised. The protein assembly was checked for invalid residues / any missing atoms and the overlapping of 25 residues at the interface were corrected using the rotamer library in Discovery studio. The dimer assembly was read with CHARMm force field, heavy atoms constrained, and energy minimized using a cascade of steepest descent 1000 steps and conjugate gradients for 1000 steps in Discovery studio^41^ and thus prepared molecules was used for docking studies.

Using prepare ligand modules, the 2D coordinates of Coomassie blue (CBBG) were converted into 3D and energy minimised using smart minimiser for 2000 steps via minimize ligands tools in Discovery studio.^42^ Gold molecular docking program CSD 2020^42^ was utilized to dock the energy minimized ligand CBBG. The NMR CSP data was used to define the active site residues involved in binding of the CBBG and default parameters and GOLD PLP fitness scoring function available with GOLD docking program was used to rank the poses.^42^

### Cytotoxicity assay

Human ovarian cancer SKOV3 cells were obtained from ATCC (ATCC, Manassas, VA, United States). It was maintained in (Roswell Park Memorial Institute) RPMI-1640 medium supplemented with 10% fetal bovine serum (FBS) and antibiotics (100 μg/mL penicillin and 100 μg/mL streptomycin). It was incubated at 37 °C in a humidified atmosphere, 5% CO_2_. The cells were sub-cultured every 2–3 days.

The cytotoxicity of the given molecule (CBBG) on SKOV3 cells was determined by MTT (3-(4, 5-dimethylthiazol-2-yl)-2, 5-diphenyl tetrazolium bromide). Cells were seeded at a density of 1 × 10^4^ cells per well in 96-well plates Each well contained 100 μl of cell suspension, and the plates were incubated for 24 h at 37 °C under 5% CO_2_ to obtain a monolayer culture. After 24 h of incubation, the medium was removed from each well. Each well was washed with PBS and then 100 μl drug solution solubilised in the incomplete medium in the concentration of 5, 10, 15, 20, 30, 40 μM was added and the experiment was performed in triplicate for each CBBG concentration. The plates were incubated for 24 h in 5% CO_2_ incubation. The supernatant was removed from each well of the plate. Then 100 μl of MTT reagent (5 mg/ml) was added and incubated for 3 h at 37 °C in the CO_2_ incubator. The MTT solution was then discarded and 100 μl of DMSO (solubilising reagent) was added. The plates were placed in a dark place for 10 minutes to solubilize the formations of purple crystal formazan. The absorbance was measured using a microplate reader at a wavelength of 570 nm. The results were used to construct a Standard Graph by taking percentage cell viability in Y-axis against a concentration of the drug in X-axis.

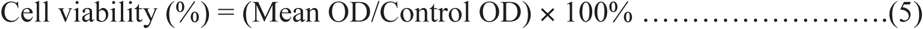

## Abbreviations

HI: Human insulin
CBBG: Coomassie Brilliant Blue G-250
CD: Circular Dichroism
NOEs: nuclear Overhauser effect
NOESY: nuclear Overhauser effect spectroscopy
TOCSY: total correlation spectroscopy
trNOESY: transferred NOESY
ThT: thioflavin T
ITC: Isothermal calorimetry
MD: molecular dynamics simulation
CSP: chemical shift perturbation
CTD: C-terminal domain
PLP: piecewise linear potential

## Author Contributions

NCM and AB envisaged the idea, designed the experiments, and analyzed the experimental data, SD and RP performed the experiments; RP analyzed the NMR data with AB; SD, RP, AB, NCM analyzed the results; SD, RP, AB, and NCM wrote the manuscript, BNR helped SD in CD data analysis; SS performing cell cytotoxity experiment in the laboratory of Snehasikta Swarnakar; AH performed molecular modeling studies, AS also helped in initial docking; AS and AH for helpful discussion and reviewing the manuscript.

## Acknowledgement

This work was partly supported by Department of Biotechnology (BT/PR29978/MED/30/2037/2018 to AB) Govt. of India and partly by Bose Institute intramural extramural research fund (R/16/19/1615 to AB). Sandip Dolui thanks CSIR network project BSC0113 for funding support. RP thanks UGC for the research fellowship. The authors thank J. Mandal and T. Murganandan for recording ITC, CD, and AFM measurements. Thanks are also the Directors and associates of the central instrumental facility of CSIR-IICB and Bose Institute for providing all of the instrument facilities.

## Notes

### Competing Interest Statement

The authors have declared no competing interest.

