## Supporting Information for "Mechanistic Studies of the Stabilization of Insulin Helical Structure by Coomassie Brilliant Blue"

*Running title: CBBG stabilizes 'partial unfold state' of insulin*

[$] both contributed equally

** Address correspondence to*

*Nakul C. Maiti, Division of Structural Biology and Bioinformatics, CSIR-Indian Institute of Chemical Biology, 4, Raja S.C. Mullick Road, Kolkata 700032, India.*

**

*Anirban Bhunia, Department of Biophysics, Bose Institute, P-1/12 CIT Scheme VII (M), Kolkata 700054, India.*

**

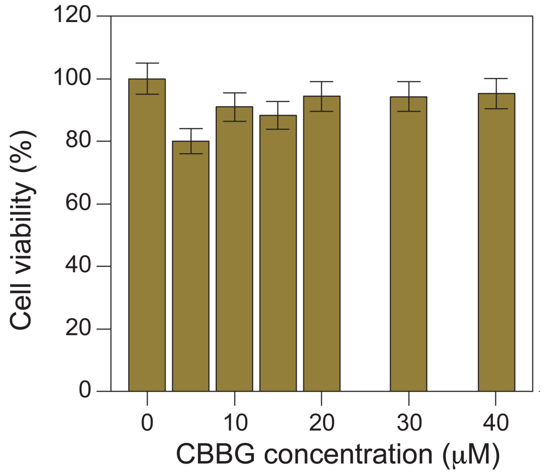


**Figure S1.** Cytotoxicity studies of the CBBG against SKOV cell line.

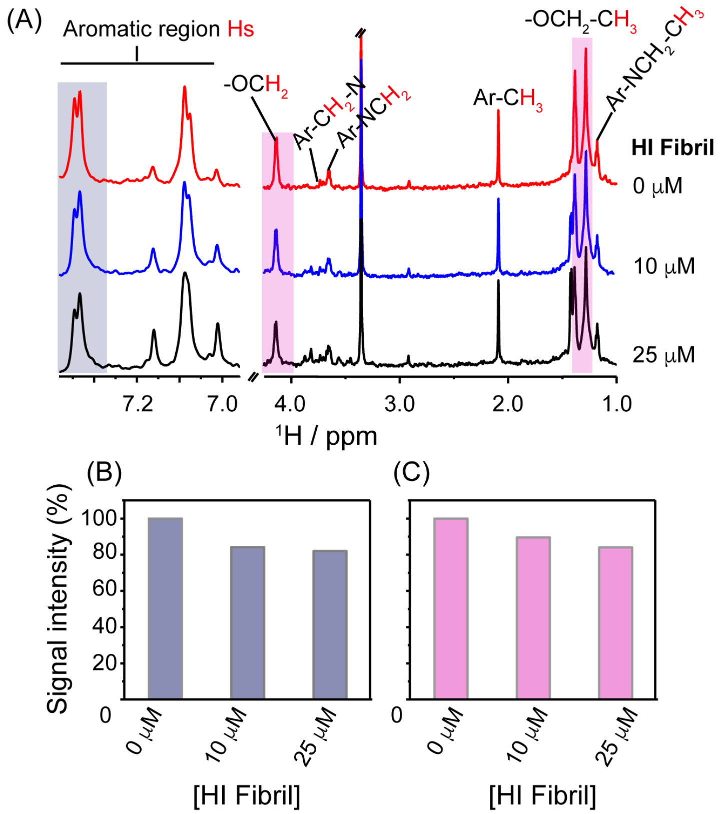


**Figure S2.** (A) Selected region of one-dimensional $^{1}$H NMR spectra of 500μM CBBG recorded at 25°C (using Bruker Avance III 700 MHz) showing the line width broadening in the aromatic and aliphatic region upon addition of insulin fibril. % of $^{1}$H NMR signal intensity were calculated from the highlighted blue (B) and purple (C) colors region as a function of insulin fibril concentration (0, 10 and 25 μM). CBBG dissolved in 25 mM HCl, 100 mM NaCl buffer with 10% $D_2O$ at pH~1.6 at 25°C.

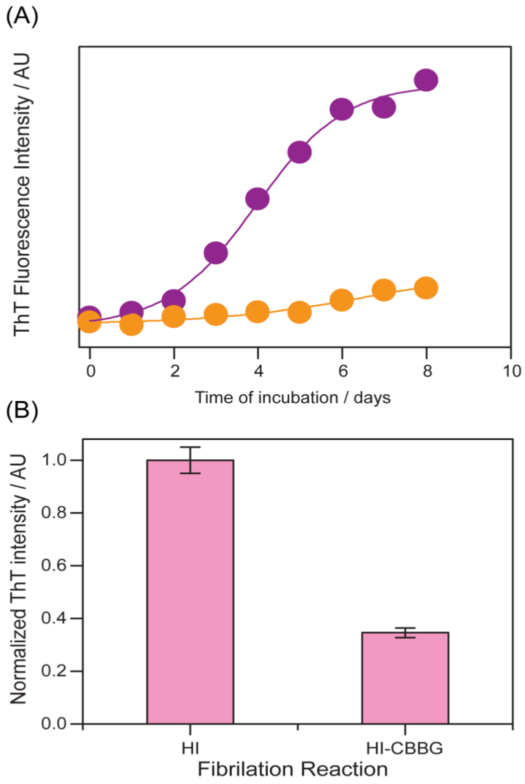


**Figure S3.** Thioflavin T fluorescence of insulin aggregation and its inhibition by CBBG (100 µM). (A) absence and presence of CBBG at insulin: CBBG ratio of 1:1 of 100 µM insulin incubated at 60 $^{o}$C, phosphate buffer, 0.1M NaCl, pH 7.2, for 8 days. The absence (violet trace) and presence CBBG (orange trace).(B) Normalized ThT fluorescence intensity of 100 µM insulin incubated in the absence and presence of CBBG.

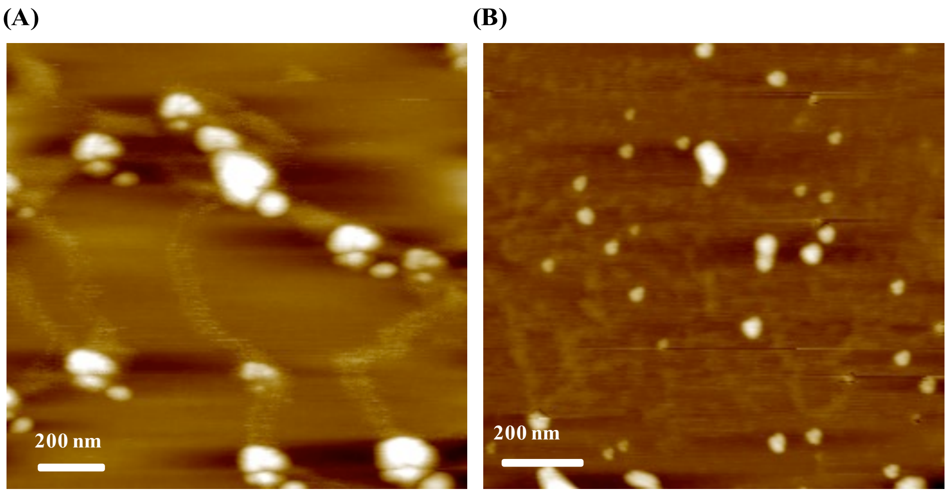


**Figure S4.** (A) AFM images of insulin aggregates after 8 days of incubation. (B) AFM image of the insulin incubation with CBBG after 8 days. (incubation condition, insulin 100 µM and CBBG 100 µM, T = 60 $^{o}$C, pH = 7.2, 100 mM NaCl).

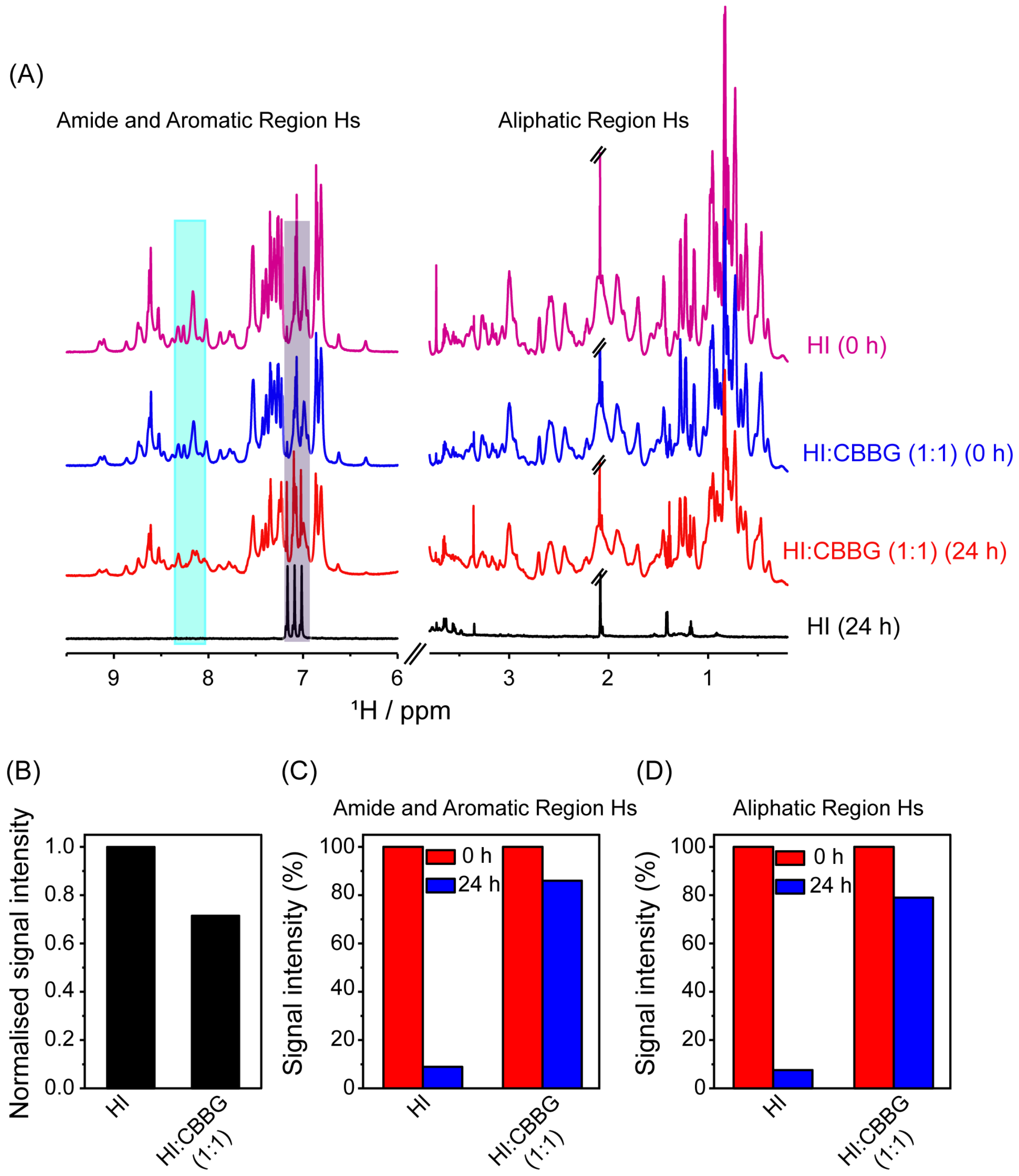


**Figure S5.** The aggregation kinetics of insulin were acquired by one dimensional $^1$H NMR. (A) Time-lapse $^1$H NMR spectra of 1mM insulin and/or insulin treated with equimolar CBBG in 10 mM sodium phosphate buffer (pH 2.0) with 10 mM NaCl. (B) Normalised signal intensity of insulin upon addition of CBBG at 1:1 molar ratio. The percentage of $^1$H NMR signal intensity of amide and aromatic region (C) and aliphatic region (D) of insulin and/or insulin+CBBG (1:1 molar ratio) for 0 and 24 h incubation at 60 $^o$C, calculated from the integrated total peak intensities of corrosponding regions. Appearence and disappearance of new peaks after heating are highlighted. All the spectra were recored on Bruker Avance III 700 MHz and at 25$^o$C.

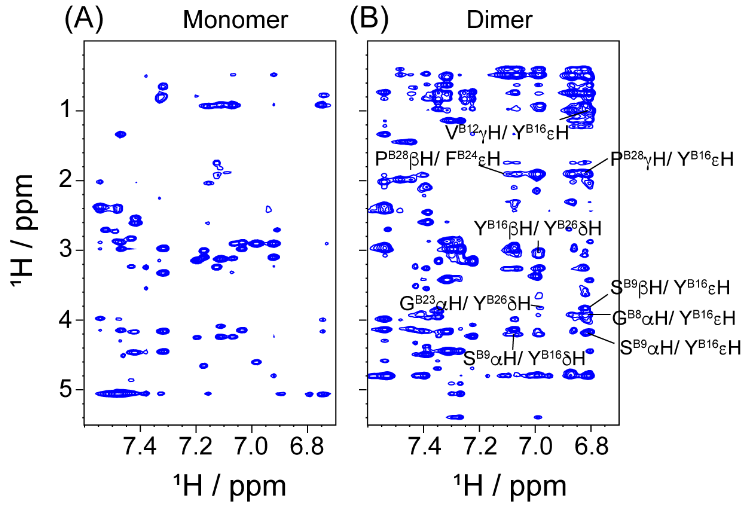


**Figure S6.** Structural difference between monomer and dimer insulin using two dimensional (2D) NMR. (A) 2D $^1$H-$^1$H NOESY spectra of insulin monomer in 20% acetic acid-d$_4$ containing 10% $D_2O$ at pH 1.9. (B) NOESY spectra of insulin dimer in 10 mM sodium phosphate buffer, 10 mM NaCl containing 10% $D_2O$ at pH 2.0. Intermonomer NOEs were detected and assigned in insulin dimer but not in monomer. NMR spectra were acquired at 25$^{o}$C on 700 MHz Bruker NMR spectrometer.

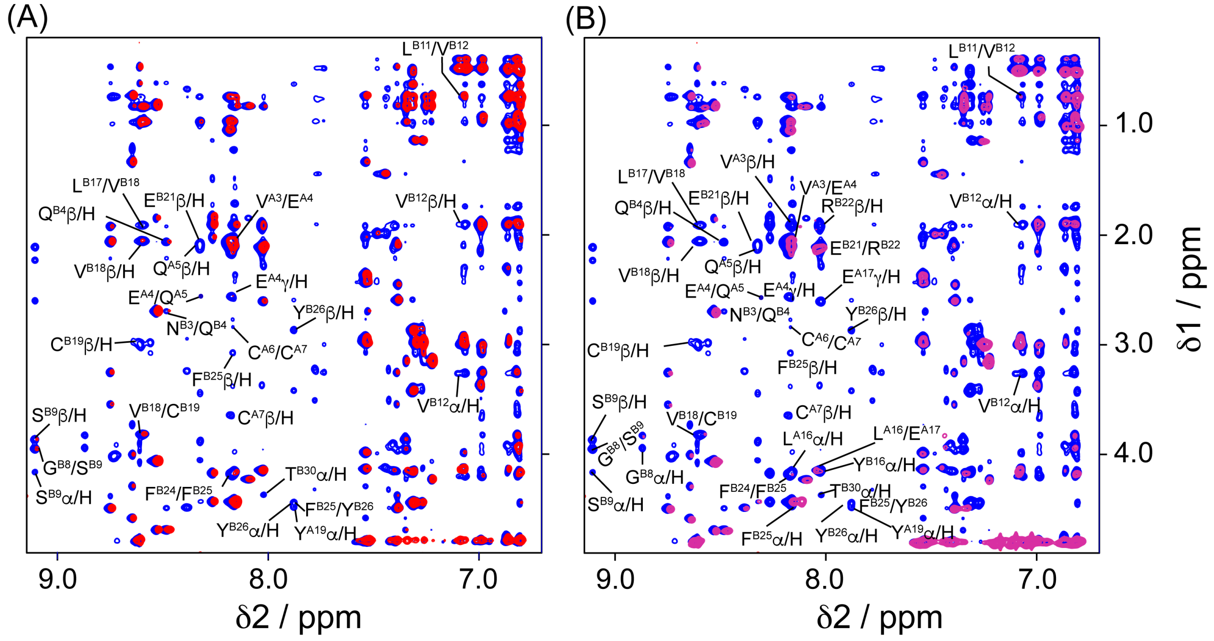


**Figure S7.** 2D homonuclear NOESY NMR spectra of 1 mM human insulin alone (blue) and insulin+ CBBG (1:1 molar ratio) before heating (red) (A) and after heating (purple) (B) at 60$^{o}$C for 24 h. NMR samples were prepared using 10 mM sodium phosphate, 10 mM NaCl buffer with 10% $D_2O$ (pH 2.0). The NMR spectra were acquired using Bruker Avance III 700 MHz NMR spectrometer at 25 $^{o}$C.

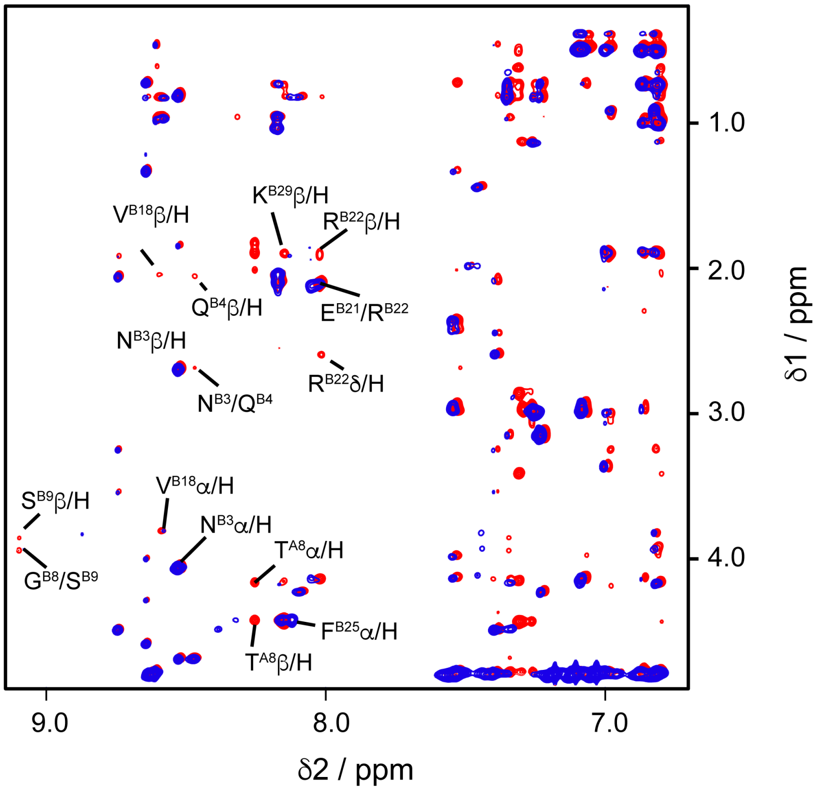


**Figure S8.** 2D $^{1}$H-$^{1}$H NOESY NMR spectra of 1 mM human insulin the presence of equimolar CBBG (1:1) before (red) and after heating (blue) for 24 h at 60 °C. The overlay NMR spectra shows peak broadening as well as chemical shift change after incubated at 60 °C in sodium phosphate buffer (pH 2) using 700 MHz Bruker NMR spectrometer.

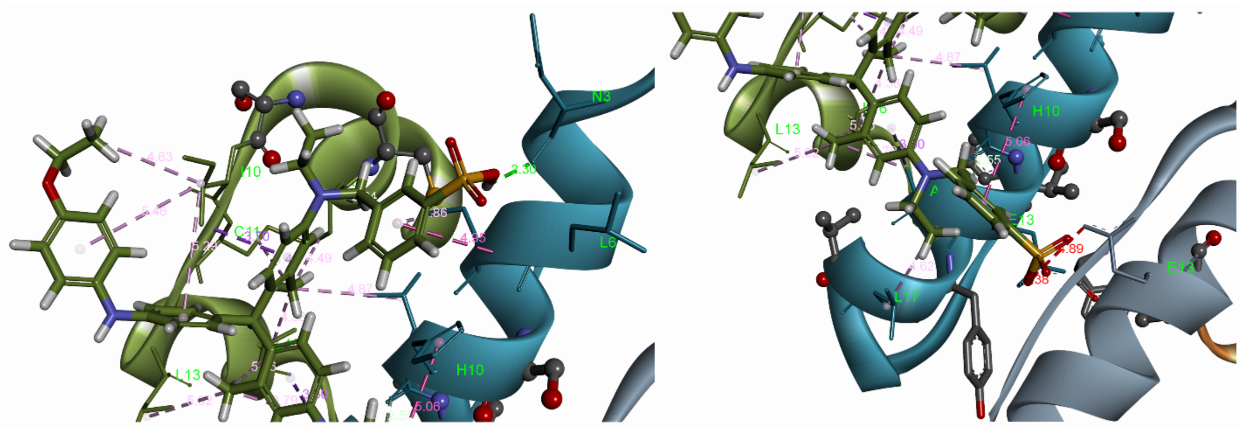


**Figure S9**, 3D Interaction sections of Trainline substitutions of CBBG, A) Interactive types and distances of Two aniline Fragments at A subunit. B) Aniline fragment with 3-sulfonate-phenyl group with B subunit.

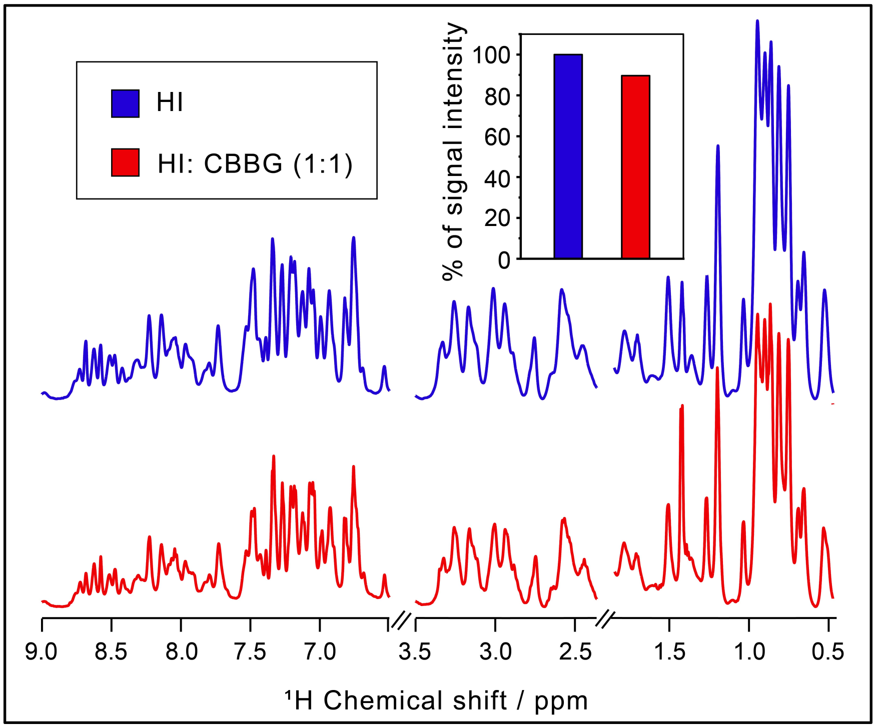


**Figure S10.** One dimensional $^{1}H$ NMR spectra of insulin before (blue) and after (red) addition of equimolar CBBG in 20% acetic acid-$d_4$ and 10% $D_2O$ at pH 1.9 (Bruker Avance III 700 MHz NMR spectrometer, 298 K). $^{1}H$ NMR signal intensity (%) of seclected region in insulin reduces to 90% from 100% upon addition of CBBG at 1:1 molar ratio.

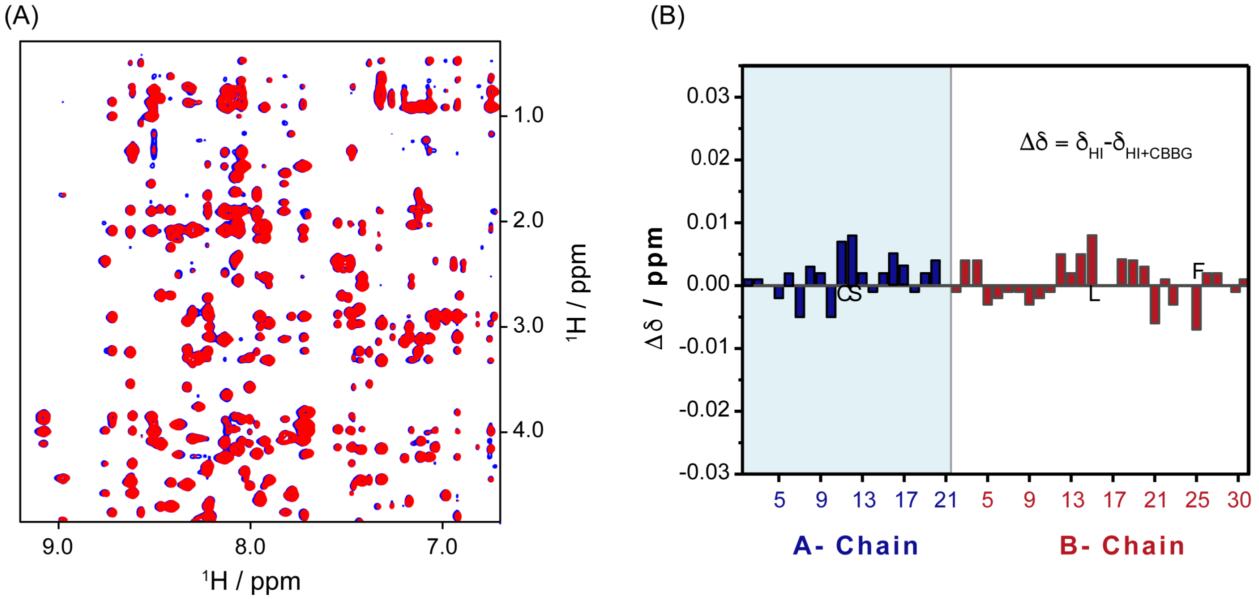


**Figure S11**. (A) 2D-NOESY NMR spectra of 350 μM human insulin in 20% acetic acid-$d_4$ + 10% $D_2O$ (pH 1.9) in the absence (blue) and presence (red) of equimolar CBBG (1:1) at 25°C using Bruker Avance III 700 MHz NMR spectrometer. (B) α-H chemical shift difference of residues from insulin A-chain (blue) and B- chain (red) between insulin alone and insulin-treated with CBBG at 25 °C.
